# CRISPR with Transcriptional Readout reveals influenza transcription is modulated by NELF and can precipitate an interferon response

**DOI:** 10.1101/2024.11.14.623683

**Authors:** Alison C. Vicary, Sydney N.Z. Jordan, Marisa Mendes, Sharmada Swaminath, Lennice K. Castro, Justin S. Porter, Kevin D. Vo, Alistair B. Russell

## Abstract

Transcription of interferons upon viral infection is critical for cell-intrinsic innate immunity. This process is influenced by many host and viral factors. To identify host factors that modulate interferon induction within cells infected by influenza A virus, we developed CRISPR with Transcriptional Readout (CRITR-seq). CRITR-seq is a method linking CRISPR guide sequence to activity at a promoter of interest. Employing this method, we find that depletion of the Negative Elongation Factor (NELF) complex increases both flu transcription and interferon expression. We find that the process of flu transcription, both in the presence and absence of viral replication, is a key contributor to interferon induction. Taken together, our findings highlight innate immune ligand concentration as a limiting factor in triggering an interferon response, identify NELF as an important interface with the flu life cycle, and validate CRITR-seq as a tool for genome-wide screens for phenotypes of gene expression.

**Significance Statement:** Nearly every cell in the human body has the ability to detect and respond to viral infection by producing interferons. The timing and magnitude of the interferon response impacts the course of disease, and both hosts and viruses have evolved mechanisms to regulate interferon induction within infected cells. It has previously been challenging to comprehensively search for regulators of interferon expression using selection-based screens. Here we developed a CRISPR screening strategy to measure the effects of gene edits on transcription at a promoter of interest. Applying this method to study interferon transcription during influenza infection, we identified an interface between human and influenza transcription machinery that modulates the viral life cycle and influences the interferon response.

## Introduction

Transcription of type I and type III interferons is one of the first host responses to viral infection in vertebrates and is critical for controlling viral spread^1,2^. Through autocrine and paracrine signaling, interferon secreted from a subset of infected cells induces transcription of a set of interferon-stimulated genes (ISGs) that establish an antiviral state in neighboring cells. Innate and adaptive immune cells also respond to interferon signaling, promoting a systemic immune response^3^. Demonstrating the importance of this pathway for effective viral clearance, mice and humans with genetic deficiencies in interferon signaling are more susceptible to severe viral disease^4,5^. However, aberrant interferon expression can also promote harmful immunopathology^1,3,6^. Therefore, the probability and magnitude of interferon induction must exist within a narrow range to support host health.

The frequency of interferon induction early in infection is shaped by the molecular interactions between host and virus within infected cells. Interferon signaling is activated when cellular receptors bind pathogen-associated molecular patterns (PAMPs) from the virus, or damage-associated molecular patterns (DAMPs) from the host cell. In influenza infection, RIG-I is the host receptor responsible for activating interferon signaling in epithelial cells^7,8^. This process is antagonized by the flu protein NS1^9^. Flu populations lacking NS1 trigger an enhanced interferon response, which is dependent on *de novo* generation of viral RNA^10^. Influenza RNA is produced by two canonical mechanisms. First, incoming viral ribonucleoproteins (vRNPs) containing the negative-sense viral genome segments are transcribed in the nucleus to generate capped and polyadenylated mRNAs. After these mRNAs are translated to produce viral proteins, newly synthesized viral polymerases replicate the genomes, generating positive-sense complementary RNP (cRNP) intermediates and new negative-sense vRNPs^11^. The primary RIG-I ligand in influenza infection is thought to be the viral RNA genomes, which have a 5’ triphosphate and a double-stranded region where the complementary 5’ and 3’ ends bind to each other^12^. Various aberrant viral RNAs have been shown to associate with enhanced interferon responses to influenza, including deletion-containing viral genomes (DelVGs) and mini viral RNAs (mvRNAs)^13–17^.

In addition to viral factors either positively or negatively influencing the interferon response, a number of different host pathways can impact its induction. Even when provided with a pure, uncomplicated, innate immune agonist such as the dsRNA mimetic poly(I:C), interferon is still produced in only a fraction of cells, implying that there may be layers of host regulation and stochasticity governing the probability of interferon production^18–20^. During infection, host processes may generate DAMPs, and interferon signaling triggers positive and negative cellular feedback loops that regulate further interferon production^21–25^. Other host factors influence viral progression through the life cycle, which is itself a highly heterogeneous process often varying in productivity by several orders of magnitude, and might therefore influence the probability of detection^26,27^. Therefore, the magnitude of the interferon response in a given viral infection is the result of a complex network of interacting factors that contribute in different contexts.

There have been very few studies performing genome-wide identification of host factors that modulate interferon induction^28,29^. The scarcity of these studies likely reflects the difficulty of assessing interferon production, a process that can be incredibly stochastic and heterogeneous^30,31^. Critically, in addition to the fact that not all cells will produce interferon during a given stimulus, even the amounts of interferon produced by a given cell in a population at a given time point may differ. This is challenging to appropriately capture when relying on methods that sort cells into simple yes/no pools based on expression. Addressing these challenges with a simplified approach, we developed a novel CRISPR screening strategy that measures the effects of different gRNAs on transcription levels from a promoter of interest, which we have termed CRISPR with Transcriptional Readout (CRITR-seq).

Using our newly-developed method, we have performed a genome-wide screen for host factors that modulate interferon transcription in response to influenza A virus infection. As expected, we identify RIG-I signaling pathway members as well as host genes supporting viral infection as required for robust interferon production. In terms of new findings, we identify that loss of the Negative Elongation Factor (NELF) complex leads to increased flu transcription, which corresponds with a dramatic increase in interferon induction, leading us to more broadly identify viral transcription as a key contributor to interferon induction by flu.

## Results

### CRITR-seq measures the effects of gRNAs on transcription at a reporter promoter

The most common form of CRISPR screen involves using CRISPR-Cas9 to generate a pool of edited cells, selecting for a phenotype of interest, and sequencing genomic DNA to identify integrated guide RNAs (gRNAs) that are enriched or depleted by selection. However, this cannot be applied to phenotypes that do not directly determine cell survival. One way to overcome this limitation is by using reporter constructs and cell sorting to apply selection to phenotypes of gene expression. While these methods are informative, they may reduce complex regulatory differences to a binary (yes/no) readout. To provide a more quantitative readout of factors that modulate gene expression, we designed a new CRISPR screen that directly measures the effects of different gRNAs on transcription levels from a promoter of interest.

Our approach, which we call <u>CRI</u>SPR with <u>T</u>ranscriptional <u>R</u>eadout (“CRITR-seq”), is a modification and combination of two prior methods. The first was developed for yeast, called CRISPR interference with barcoded expression reporter sequencing (CiBER-seq)^35^. CiBER-seq uses a barcode embedded downstream of a reporter promoter. After sequencing is used to link a barcode to the appropriate guide, transcribed barcodes are used to measure guide impacts on transcription at the reporter promoter. While this method is useful in yeast, the lentiviral constructs generally used for human CRISPR screens can recombine^36^, making it difficult to link a barcode to a guide. To overcome this limitation, we incorporated the entire gRNA expression construct downstream of our reporter promoter, such that the guide would be present within the reporter RNA. We took inspiration for this approach from CRISPR droplet sequencing (CROP-seq), which uses a similar design to recover guide RNA sequences from integrated lentiviruses in single-cell RNA-seq^37^. Our design is explained in Fig. 1A-B.

**Figure 1.**
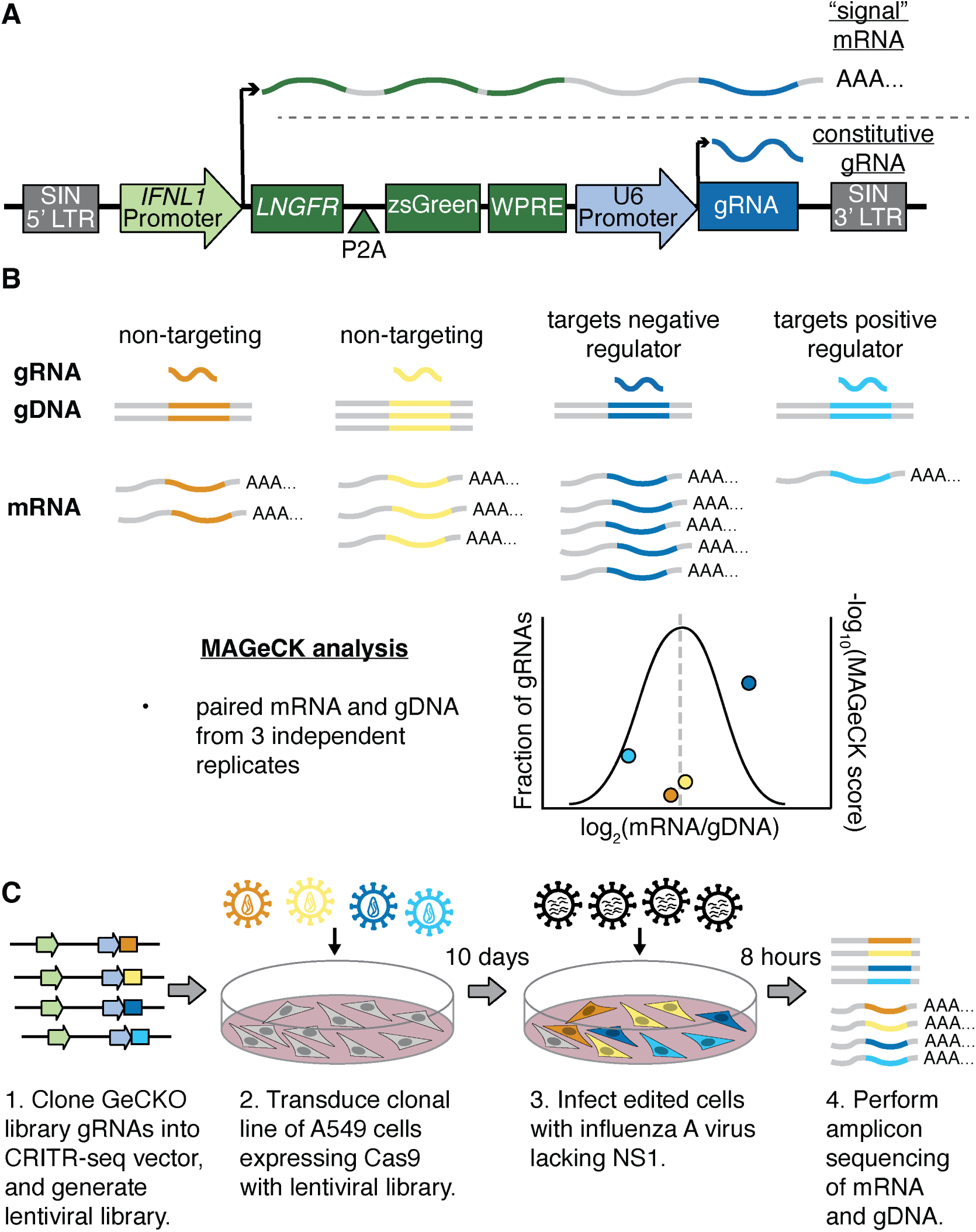
CRISPR with Transcriptional Readout (CRITR-seq) workflow. (A) CRITR-seq vector. gRNA is constitutively transcribed from the U6 promoter. gRNA sequence is also present in the reporter mRNA transcribed from the *IFNL1* promoter, serving as a barcode indicating which edit occured in the transcribing cell. SIN LTR = self-inactivating long terminal repeat, *LNGFR* = low-affinity nerve growth factor receptor, WPRE = woodchuck hepatitis virus posttranscriptional regulatory element. (B) Model analysis of a CRITR-seq screen. gRNA sequence is measured in both genomic DNA and reporter mRNA. The mRNA/gDNA ratio for each gRNA sequence represents the normalized *IFNL1* transcription levels for that guide. Graph represents a distribution of all gRNAs based on their mRNA/gDNA ratio, with individual examples highlighted as colored points plotted based on their hypothetical MAGeCK^38^ score. (C) Workflow for the primary CRITR-seq screen in this study. 3 libraries were generated independently from initial cloning of gRNA sequence from the GeCKO library^39^. A clonal line of Cas9-expressing A549 cells was transduced with lentivirus carrying the CRITR-seq vector at an MOI of 1.5, with an assumption that epistasis or conflict would be rare. After allowing 10 days for gene editing, edited cells were infected with NS1_mut_ influenza A virus at an MOI of 2.

In this study, we used CRITR-seq to perform a genome-wide CRISPR knockout screen for factors that modulate interferon induction during influenza A virus infection. For this purpose, we used the promoter for the type III interferon *IFNL1*. In A549 cells, a lung epithelial carcinoma cell line, expression of type I and type III interferons are highly correlated, and we have previously validated that our reporter promoter correlates well with the transcriptional response at the endogenous promoter in this cell line^40,41^. We use a type III rather than a type I interferon promoter as in these cells, during flu infection, we have found the former to drive a stronger response than the latter^40^. In each cell containing the CRITR-seq construct, both a prototypical gRNA is transcribed for Cas9-editing, and an additional, longer, reporter mRNA is transcribed whenever the interferon promoter is active (Fig. 1A).

After generating a pool of edited cells, the abundance of each gRNA sequence in the longer reporter mRNA should correlate with the effects of that guide on interferon transcription (Fig. 1B). To account for the frequency of lentiviral integration, we normalize reporter gRNA sequence to the corresponding number of reads in the genomic DNA (gDNA). We used Model-based Analysis of Genome-wide CRISPR/Cas9 Knockout (MAGeCK)^38^ to calculate a statistical score for the enrichment or depletion of gRNAs targeting each gene, based on a null distribution drawn from non-targeting guides. We expect that gRNAs targeting positive regulators of interferon expression will be depleted in the mRNA, while gRNAs targeting negative regulators will be enriched.

### Analysis of depleted guides confirms efficacy of CRITR-Seq

The workflow for the primary CRITR-seq screen in this study is shown in Fig. 1C. This screen used a strain of influenza that lacks the innate immune antagonist NS1, to enhance our ability to detect interferon-producing cells. An additional feature of this virus is that segments 1-7 are derived from the lab adapted strain A/WSN/1933, whereas the eighth segment, where we have mutated NS1, is from A/PR8/1934, as, for reasons that remain unclear to us, this virus grows better than its WSN counterpart. In addition to our large screen, we also performed a smaller screen consisting of 216 genes targeted by three guides each with 100 control non-targeting guides to assess the reproducibility of our approach. Design of this library is described in Fig. S1.

Our primary screen maintained high levels of guide diversity, with well over 90% of guides observed in both DNA and RNA samples Fig. S2. We anticipated most guides would have no effect on transcription of our reporter — if so the number of times a guide is present in our DNA samples should match its abundance in the RNA samples for the majority of guides, which we observed with Pearson correlation coefficients approaching 0.9 (Fig. S3). Inter-replicate correlations were also high for both DNA and RNA measurements. Similar correlation coefficients were observed at the gene-level as for individual guides (Fig. S4). Lastly, from our smaller repeat CRITR-seq experiment we observed that, of the top 10% of enriched and top 10% of depleted genes in that dataset, there was agreement in either enrichment or depletion ∼90% of the time with our larger dataset (Figs. S5 and S6). Therefore for genes where we have statistical confidence there is a high degree of replicability between completely independently performed experiments.

Moving to predicted effects on interferon induction by knockout of a given gene, we first present the distribution of MAGeCK scores versus log-fold change (Fig. 2A). We would anticipate that, broadly, two features should be absolutely required for an interferon response, an intact RIG-I signaling pathway, and the ability to be infected by influenza virus (Fig. 2B,C). Regarding the former, our results show that edits targeting genes required for RIG-I signaling are largely predicted to have a negative effect on interferon induction, as anticipated (Fig. 2B). While edits targeting *RIG-I* are oddly predicted to enhance interferon transcription, we provide a potential explanation in Fig. S7. Critically, owing to a single strong effect guide, *RIG-I* was still ranked in the top 10% of depleted genes by MAGeCK scoring (Fig. 2D). Similarly guides associated with edits anticipated to block flu entry are also depleted in our screen, matching expectations (Fig. 2C).

**Figure 2.**
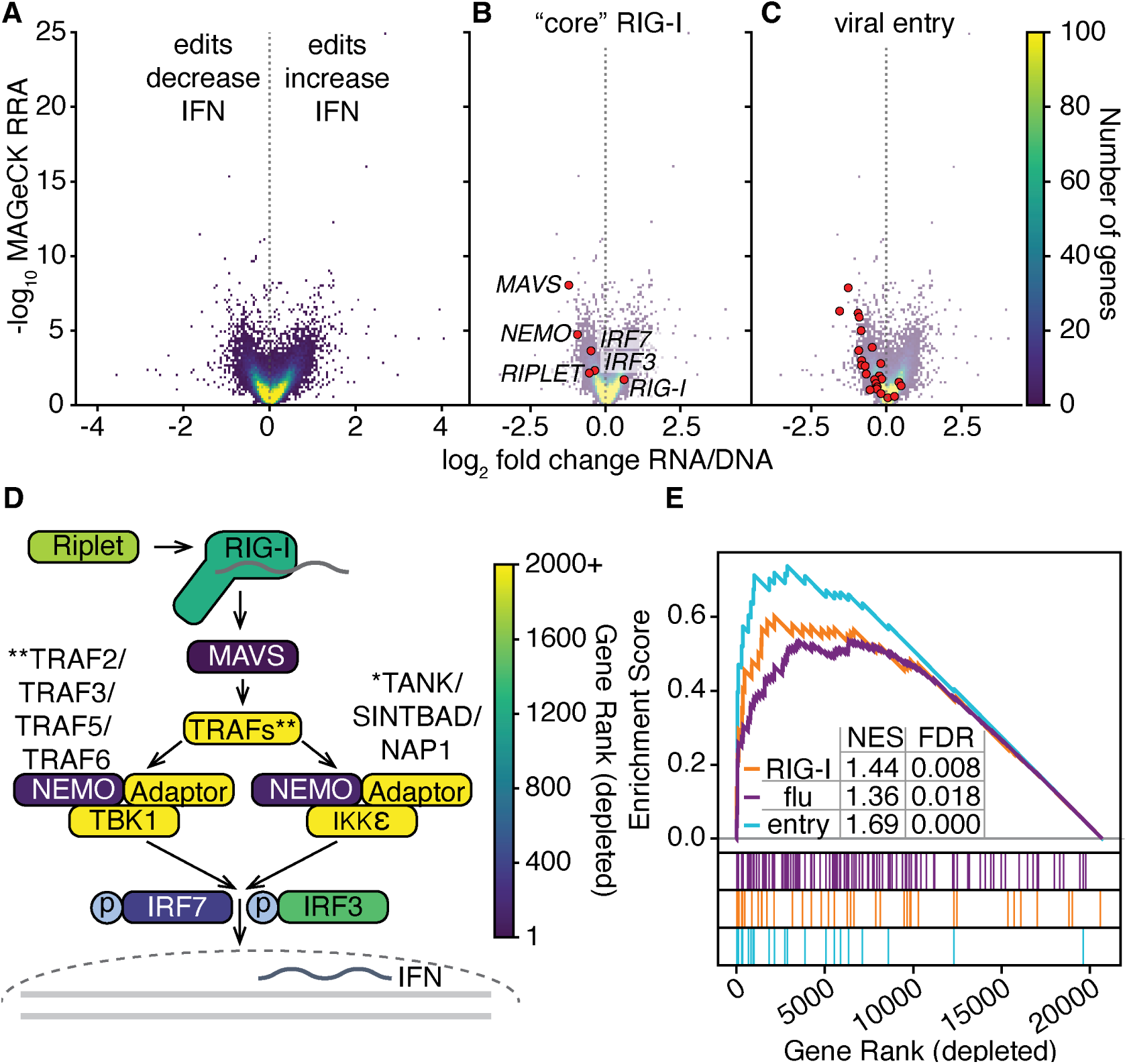
Validation of CRITR-seq through recovery of factors required for infection and RIG-I signaling (A) MAGeCK RRA (robust ranking aggregation) scores and predicted log_2_ fold change of reporter expression (RNA) relative to integration events (DNA) as calculated across three independent replicates of CRITR-seq performed in A549 cells infected with NS1_mut_ virus at an MOI of 2, and analyzed 8 hours post infection. Each individual pixel represents a single gene, overlapping genes shown by heatmap. All calculations relative to internal non-targeting controls. Y-axis is the RRA associated with a one-tailed test of increased (right of dotted line) or decreased (left of dotted line) guide abundance in the RNA pool. Full data provided as Table S1. (B) Plot as in A with core RIG-I genes noted, those that are either non-redundant, or at least partially non-redundant in the case of *IRF3* and *IRF7*. (C) Plot as in A with genes annotated as likely required for viral entry as noted in a prior CRISPR screen by Li *et al*.^42^ (D) RIG-I signaling pathway. Genes colored by MAGeCK ranking of depleted guides. All genes ranked 2000 and above colored identically. Despite apparent enrichment of *RIG-I* a single strong-effect guide leads to its prediction as required for interferon signaling. (E) Gene-set enrichment analysis of the indicated pathways, either using genes required for flu infection as identified by Li *et al*., or the RIG-I pathway (GOBP RIG I SIGNALING PATHWAY) from the gene ontology consortium.^43–45^ Gene lists provided in Table S3. NES, normalized enrichment score, FDR, false discovery rate.

To quantitatively characterize these associations, we performed gene set enrichment analysis (GSEA), for a larger set of RIG-I associated genes (the initial core set is too small for GSEA), and those genes identified as required for infection by a prior CRISPR screen (Fig. 2E)^42–45^. As an additional test, we used a subset of those genes required for flu entry as shown in Fig 2C, as while not all steps required for flu productive infection may impact interferon production, entry is an unambiguously required step. We see for all three gene sets there is substantial enrichment of the indicated genes in the set of those predicted to be required for interferon induction. Comparing our two flu gene sets, we see an even stronger enrichment of those genes required for viral entry alone.

Lastly, to verify our method we chose to examine our top depleted gene with a predicted >2-fold effect size, *CSNK2B*, which encodes the regulatory subunit of casein kinase 2 (CK2). CK2 was recently found to phosphorylate CMTR1, required for both flu efficient flu replication as well as the interferon response^46^. Consistent with this work, and verifying our results, we find that silencing *CSNK2B* reduces interferon production from the endogenous type I interferon locus during infection with NS1_mut_ influenza (Fig. S8).

### Loss of the Negative Elongation Factor complex enhances interferon induction by flu

We next examined the genes with guides enriched in the mRNA, indicative of suppressors of the interferon response. The 4^th^ through the 6^th^ top scoring enriched hits, *PKRA*, *TADA1*, and *NMI*, either directly suppress an interferon response, are components of a complex observed to suppress the interferon response, or mediate an antiviral response against influenza and thus may reduce replication and ligands available for RIG-I.^47–49^ The top three hits in our entire screen are all components of the Negative Elongation Factor (NELF) complex: *NELFB*, *NELFA*, and *NELFCD* (Fig. 3A), none of which have any defined roles in interferon transcription or flu infection. The NELF complex consists of NELF-A, NELF-B, either NELF-C or NELF-D, and NELF-E. *NELFE* was also recovered as suppressing interferon production, as the top 26^th^ hit enriched in our screen. Our smaller screen included *NELFB*, *NELFA*, and *NELFCD* (but not *NELFE*), and repeated the enrichment observed in our larger CRITR-seq experiments (Fig. S9).

**Figure 3.**
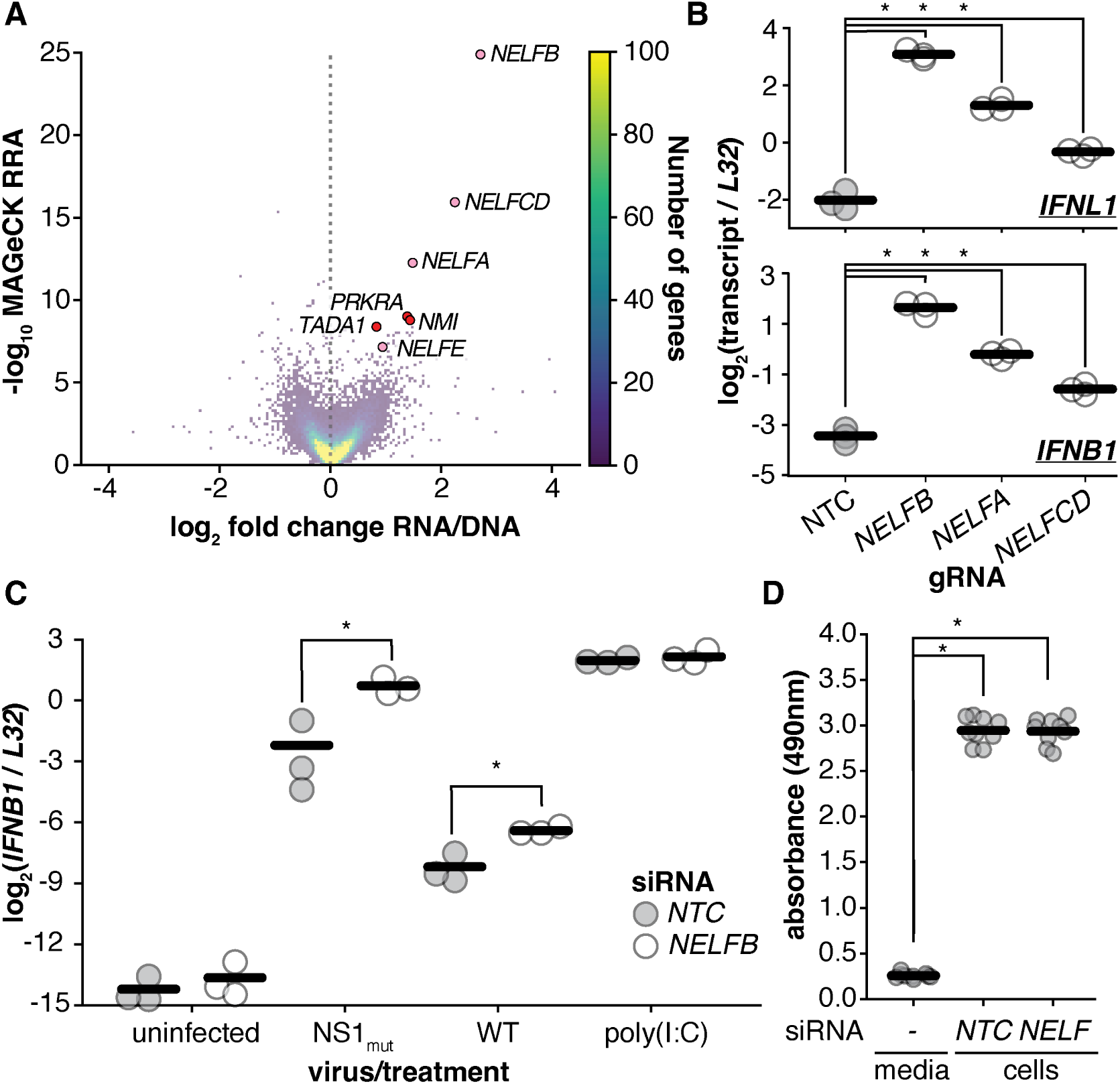
Loss of the Negative Elongation Factor complex enhances interferon induction by flu. (A) Same distribution of genes from CRITR-seq screen as in Fig. 2A, highighting the top six ranked enriched gentes by MAGeCK (red, pink), and genes encoding components of the NELF complex (pink only). (B) A549 cells were transfected with Cas9-gRNA ribonucleoprotein complexes with gRNAs targeting the indicated genes. 10 days after transfection, cells were infected with NS1_mut_ at an MOI of 2. RNA was harvested at 8 hours post-infection for qPCR analysis of *IFNB1* and *IFNL1* transcripts normalized to the housekeeping ribosomal gene, *L32*. NTC = non-targeting control. Asterisks indicate significant increases in expression over control, one-tailed Student’s t-test, p<0.05, n=3 with Benjamini-Hochberg multiple hypothesis correction. (C) A549 cells were treated with siRNA for 9 days and then infected with the indicated WSN strain at an MOI of 2, left untreated, or transfected with 50ng poly(I:C). RNA was harvested 8h post-treatment for qPCR analysis as in B, statistics and replicate numbers also as in B. Validation of *NELFB* knockdown is shown in Fig. S10. (D) Viability of cells treated as in C as measured by reduction of MTS reagent. t Both samples tested had significant increases in activity over media only control, neither differed significantly from one another. One-way ANOVA with post hoc Tukey, n=9, p<0.05. For B-D each point represents a single replicate, lines represent mean values.

The NELF complex regulates RNA polymerase II transcription, and is involved in the promoter-proximal pause, an early even that occurs ∼20-60 nucleotides downstream of the transcription start site^50–54^. This step is a checkpoint between transcription initiation and elongation^55–58^. This pause before productive elongation is the stage at which capping enzymes add an m^7^G-cap to the 5’ end of the nascent mRNA, which is important for mRNA stability, export from the nucleus, and translation^59–61^. In the absence of NELF, RNA polymerase II does not pause at the promoter-proximal region, but instead stalls further downstream, near the location of the +1 nucleosome^57,62^.

Given the presence of three members of the NELF complex as our top three enriched hits, we wished to confirm our findings. Using single CRISPR guides with ribonucleoprotein transfection, we targeted *NELFA*, *NELFB*, and *NELFCD* and measured type I and III interferon transcription during infection with NS1_mut_. Each gene we tested matched our predictions from CRITR-seq (Fig. 3B). The same effects were observed for type I interferon transcription, demonstrating that any impacts of NELF cannot be limited to the *IFNL1* locus and likely globally impact interferon production during flu infection.

As we validated our three NELF targets, and each is co-dependent, we chose to focus on one, *NELFB*, for further characterization.^62,63^. Because it is possible for CRISPR edits, which generally knockout a gene, to generate truncated proteins that may act as hypo- or hypermorphs, we decided to use siRNA-mediated knockdown of *NELFB* for further experiments. Consistent with the CRISPR-Cas9 results, knockdown of *NELFB* increased interferon transcription in response to both NS1_mut_ and wild-type WSN influenza (Figs. 3C). This effect appears specific to flu infection, as when we challenge cells by transfection with a dsRNA mimetic, poly(I:C), we do not observe an increase in interferon transcription.

Validating our results, we confirmed depletion of NELFB protein, that *NELFB* silencing had no marked effects on metabolic activity, that silencing itself did not produce a measurable interferon response as additionally measured by qPCR against an interferon-stimulated gene, that our silencing phenotype was complementable by overexpression of a silencing-resistant allele of *NELFB*, that the same phenotype could be observed in an alternate cell line (HEK293T), and that the effect we observed was not unique to a flu strain bearing segment 8 from WSN rather than the PR8 strain we used in our experiments (Figs. 3D, S10, S11, S12, and S13).

### Loss of NELF increases canonical flu transcription

As NELF depletion does not affect the interferon response to poly(I:C), we hypothesized that it may act by influencing some stage of the flu life cycle, increasing viral production of RIG-I ligands. Supporting this hypothesis we found that knockdown of *NELFB* led to an increase in RNA encoding the flu gene *HA*, as did Cas9-editing of *NELFB*, *NELFA*, or *NELFCD* (Figs. 4A and S14). We hypothesized that the increases we observed may be linked to flu transcription. Influenza transcription requires the viral polymerase to bind actively-transcribing host RNA polymerase II, whereupon it binds to the mRNA cap structure and uses an endonuclease to “snatch” the nascent transcript to use as a primer^64–69^. This interaction is thought to occur during the promoter-proximal pause of RNA polymerase II^64,70,71^. Given that NELF depletion affects RNA polymerase II promotal-proximal pausing dynamics^57,62,72^, we hypothesized that loss of NELF could also affect flu cap-snatching and transcription in an infected cell. Consistent with this model, which would anticipate that the effects of NELF both in enhancing viral RNA production and interferon would be unique to nuclear cap-snatching viruses, when we infected with a cytoplasmic RNA virus that does not rely on cap snatching, human coronavirus OC43, *NELFB* silencing does not enhance viral replication nor interferon production (Fig S15).

**Figure 4.**
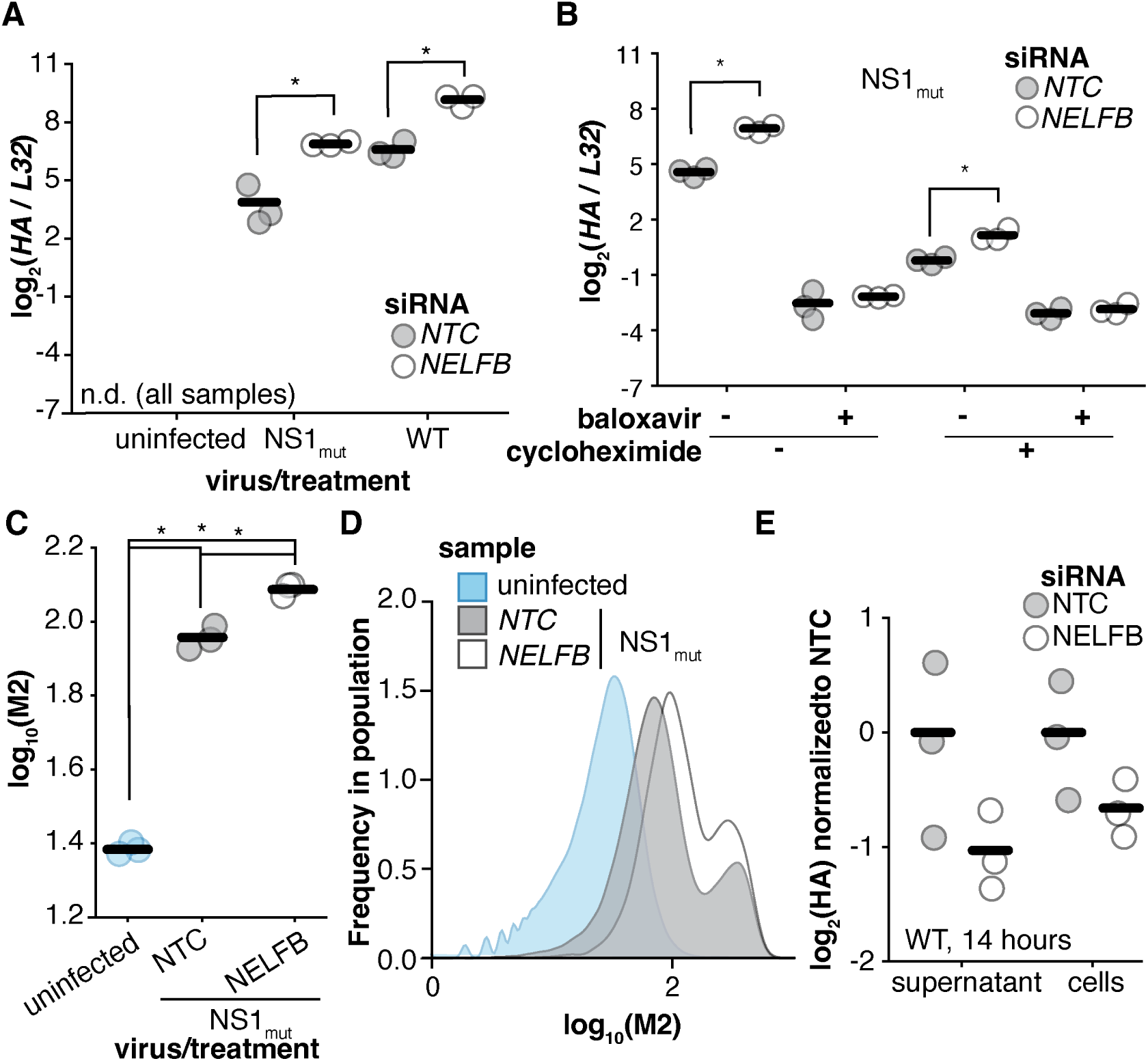
Loss of NELF increases productive flu transcription. (A) RNA from experiment performed in Fig. 3C, with *HA* transcripts measured by qPCR. n.d. = not detectable. Measurements were also made for poly(I:C) treatment and showed undetectable levels of *HA*, as anticipated. Asterisks represent significant differences between NTC and *NELFB* silencing, two-tailed Student’s t-test with Benjamini-Hochberg multiple hypothesis correction, p<0.05, n=3. (B) Cells were silenced as in A, and infected with NS1_mut_ at an MOI of 1 for 8h before RNA was harvested for qPCR analysis of flu transcript levels normalized to the housekeeping control *L32*. A549 cells were treated, or not, at the time of infection with 100 nM baloxaviric acid to suppress transcription, and cycloheximide at a concentration of 50 μg/ml to suppress replication. Asterisks indicate significant increases upon *NELFB* silencing, one-tailed Student’s t-test with Benjamini-Hochberg multiple hypothesis correction, p<0.05 n=3. *NELFB* silencing confirmed in Fig. S16. (C) A549 *IFNL1* reporter cells were silenced and infected with NS1_mut_, or not as indicated, as in A and B, except at 13h cells were stained for the viral protein M2 and analyzed by flow cytometry. Asterisks indicate significant differences, one-way ANOVA with post hoc Tukey test, p<0.05, n=3. Full flow data shown in Fig. S18. (D) M2 staining distributions of a single sample representative each from C. (E) A549 cells were treated with siRNA 9 days before infection with WT WSN at an infectous MOI of 5. Media was exchanged 2 hours post infection. 14 hours post infection, viral supernatant was collected and cells were lysed for RNA extraction. Reverse transcription was performed using universal influenza primers for the RNA from the supernatant, and random hexamer primers for the RNA from the cell lysate. No significant increase in *HA* RNA was observed, one-tailed Student’s t-test with Benjamini-Hochberg multiple hypothesis correction, p<0.05 n=3. Further qPCR analysis is shown in Fig. S20. For A-C, and E, individual points represent biological replicates, and lines represent mean values.

To test whether NELF can influence flu transcription, we measured the effects of *NELFB* depletion during cycloheximide treatment. Cycloheximide prevents translation, blocking viral genome replication but still allowing initial viral transcription. Under these conditions *NELFB* knockdown still increased *HA* RNA levels (Fig. 4B), consistent with the loss of NELF enhancing viral transcription. We next measured whether blocking viral replication and transcription, concurrently, prevents the increase in flu RNA observed during *NELFB* silencing. When we added baloxavir, a specific endonuclease inhibitor that inhibits flu cap-snatching and thus viral transcription, silencing *NELFB* no longer increased flu RNA (Fig. 4B)^73^. We therefore conclude that the increased flu RNA upon NELF depletion is dependent on transcription. To confirm this effect independent of cycloheximide treatment, we used a virus that cannot engage in genome replication due to a large internal deletion in the polymerase-encoding segment, which we also observed exhibits an increase in flu RNA upon *NELFB* silencing (Fig. S17).

While viral transcription increases during NELF depletion, we did not know if these new mRNAs are functional viral mRNAs. Using flow cytometry, we determined that knockdown of *NELFB* led to higher levels of the flu protein M2 (Fig. 4C,D, and S18), indicating that at least some of the additional flu RNA generated during NELF depletion must be supporting normal translation. We further confirmed that normal flu transcription appears enhanced by measuring the length of cap-snatched host mRNA sequence, which was unaltered by depletion of *NELFB* (Fig. S19).

Next, we tested whether NELF recruitment of the cap-binding complex (CBC) could be an explanation for the enhanced influenza transcription we observe. Shortly after 5’ caps are added to nascent mRNA, these caps are bound by the cap-binding complex (CBC), a heterodimer of nuclear cap binding protein (NCBP) 1 and 2, which could compete with flu polymerase for binding to host mRNA cap structures^74,75^. NELF depletion leads to a general decrease in NCBP1 occupancy on chromatin^61–63,76^. Therefore, we hypothesized that flu transcription increases in the absence of NELF could be due to increased availability of capped nascent mRNAs not bound by CBC. To determine the impact of CBC presence on flu transcription, we treated cells with siRNA targeting *NCBP1* and then infected them with flu that cannot engage in genome replication. *NCBP1* knockdown led to increased levels of *HA* RNA for this primary transcription-only virus (Fig. S17C), demonstrating that decreased levels of the cap-binding complex increase influenza transcription, although we cannot be certain this is the mechanism through which NELF depletion influences flu transcription without additional study.

Finally, we wanted to determine how increased transcription upon NELF depletion affects productive flu replication. We measured viral RNA levels in both cells and supernatant 14 hours post infection by wild-type virus. This time point should allow us to observe any net effect on production of viral progeny, with reduced influence from autocrine and paracrine interferon signaling. We found that after 14 hours of infection, we do not find a significant difference in flu RNA levels on *NELFB* knockdown (Fig. 4E, S20). Given previous findings that flu transcripts can make up over half the total cellular transcriptome later in infection^26^, it would make sense that flu transcription is initially accelerated upon NELF depletion but is eventually limited by other factors, reaching a similar saturation point. Knockdown of *NELFB* also did not significantly increase viral titer in the supernatant, as measured by qPCR, showing that the increased transcription early in infection does not appear to lead to similar increases in final progeny.

### Loss of NELF increases the generation of RIG-I ligands by flu

When NELF is depleted, the increased levels of viral transcription likely lead to a subsequent increase in viral replication due to the more rapid accumulation of viral proteins required for genome replication. Our observation that *NELFB* knockdown increases viral *HA* levels by a greater degree in the absence of cycloheximide than in its presence (Fig. 4B) further suggests that NELF depletion increases the generation of both flu transcripts and, as a consequence, genomic vRNA. With this synergistic increase in flu RNAs, we hypothesized that the large increase in interferon induction when NELF is depleted is due to increased production of immunostimulatory ligands.

To determine whether viral RNAs generated through transcription and replication are responsible for the enhanced interferon induction upon loss of NELF, we knocked down *NELFB* and infected cells with NS1_mut_ influenza in the presence or absence of baloxavir. In this experiment, the baloxavir treatment suppressed 99.8% of the increase in interferon induction caused by *NELFB* knockdown (Fig. 5A), suggesting that enhanced interferon caused by loss of NELF is largely dependent on *de novo* viral RNA generated through transcription and/or replication.

**Figure 5.**
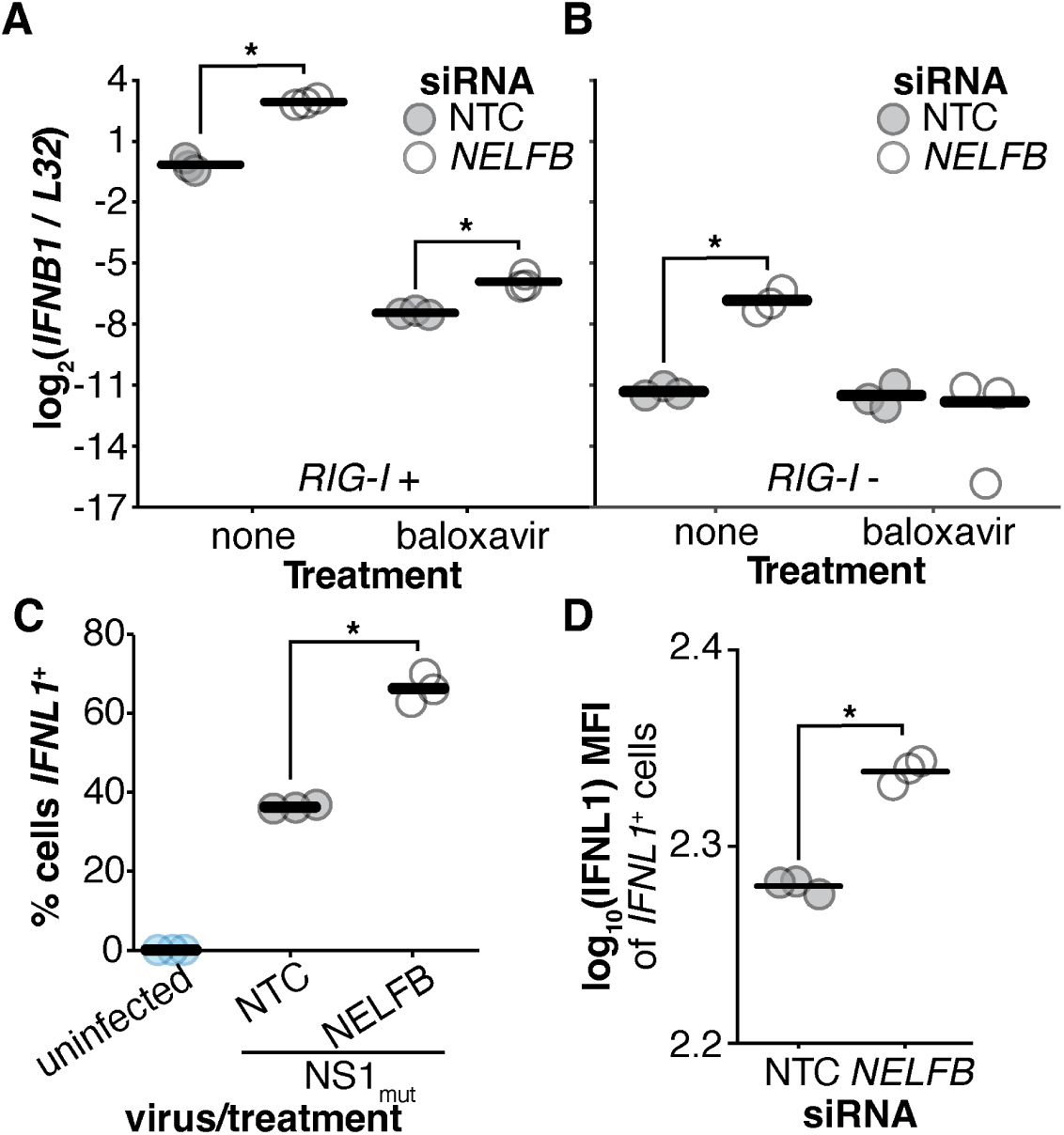
Loss of NELF increases the generation of RIG-I ligands by flu. (A-B) Wild-type A549 cells (A) or a *RIG-I* –knockout cell line derived from a single cell clone (B) were treated with siRNA for 9 days. Cells were infected with NS1_mut_ at a genome-corrected MOI of 1, with or without 100 nM baloxaviric acid added at time of infection. RNA was harvested 8 hours post infection for qPCR analysis. This *RIG-I* –knockout cell line has some residual RIG-I expression observed by Western blot after flu infection (Fig. S21). *HA* qPCR measurements in these cells are shown in Fig. S22A. (C-D) Flow cytometry experiment from Fig. 4C, with zsGreen *IFNL1* reporter results shown. (C): %*IFNL1* ^+^ cells determined as the percent of cells positive for zsGreen expression. (D) mean fluorescence intensity (MFI) of zsGreen for *IFNL1* ^+^ cells. Full flow data shown in Fig. S18. Points represent biological replicates, with lines indicating the means. Two-tailed t-tests were performed comparing the *NELFB* siRNA samples with the non-targeting control for each treatment condition, n=3. * indicates p<0.05 after Benjamini-Hochberg multiple hypothesis correction.

We repeated this experiment in a *RIG-I* –knockout A549 cell line generated in our lab (Fig. S21). In these cells, *NELFB* knockdown led to only 0.12% of the increase in interferon expression compared to wild-type cells (Fig. 5B, S21, S22). This indicates that the vast majority of enhanced interferon induction upon NELF depletion was RIG-I-dependent, which is the pathway we would anticipate to be triggered by increased flu RNA. The remaining response could be explained by small amounts of residual RIG-I, or alternative, less active, pathways.

We expect that increased concentration of immunostimulatory ligands increases the probability of receptor-ligand interactions, leading to earlier and more frequent activation of the interferon pathway. Consistent with this model, flow cytometry performed on infected cells with an *IFNL1* zsGreen reporter showed that *NELFB* knockdown led to both a greater fraction of interferon-expressing cells and increased interferon expression per cell (Figs. 5C-D).

### Transcription of deletion-containing flu genomes can contribute to interferon induction

We have seen that upon NELF depletion, the majority of the enhanced interferon induction by NS1_mut_ influenza can be attributed directly or indirectly to increased viral transcription. As multiple types of viral RNAs, including both transcripts and genomes, are increased in this context, we wanted to determine whether transcription itself participates in interferon induction apart from its role in promoting genome replication. We first tested whether, when treated with cycloheximide, silencing of *NELFB* still increases interferon production. With our NS1 knockout virus, we were unable to detect an increase in interferon transcription during cycloheximide treatment, indicating that enhanced genome replication upon NELFB depletion was likely the predominant driver of the interferon response (Fig. 6A).

**Figure 6.**
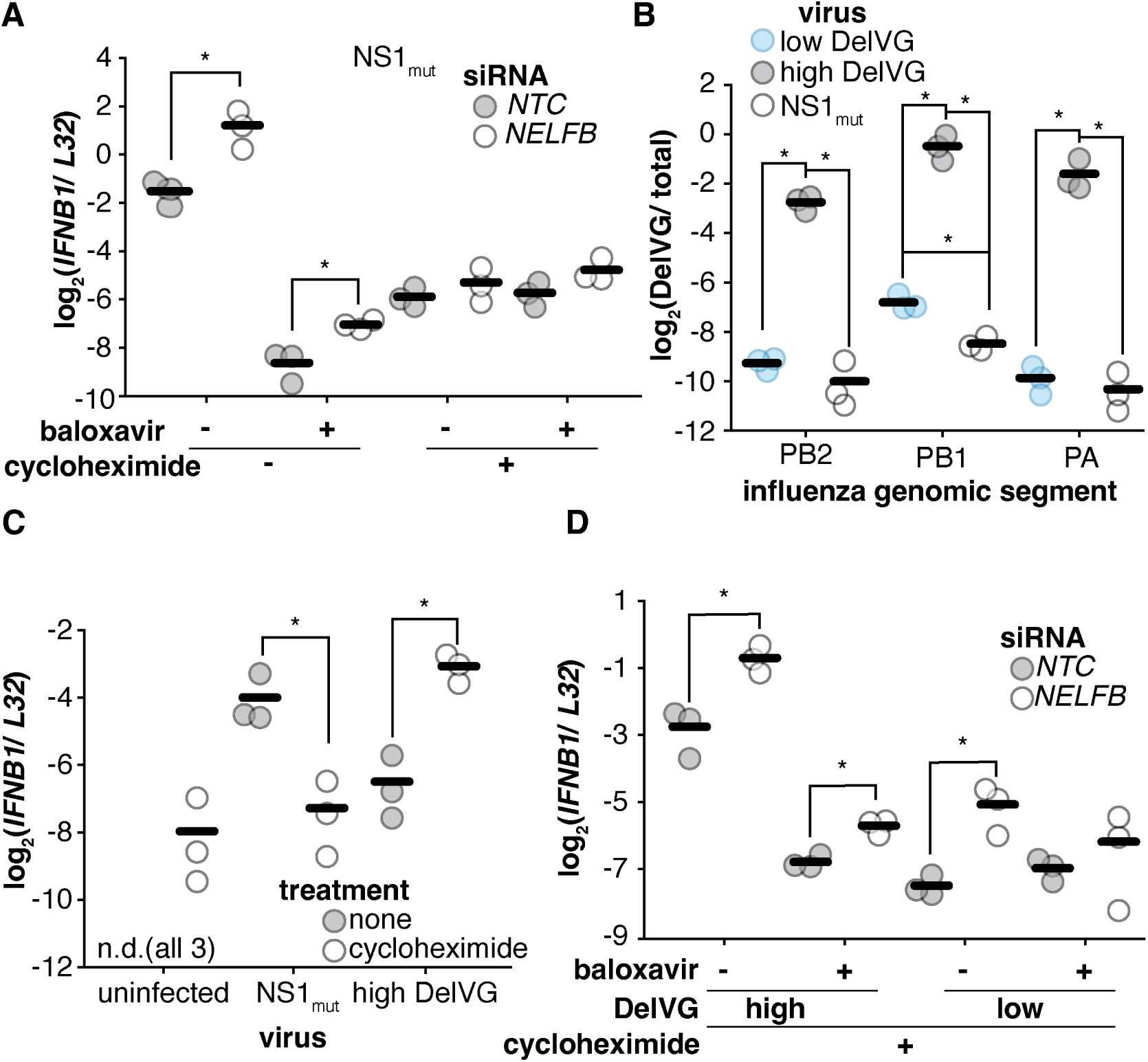
Influenza transcription contributes to interferon induction in the absence of viral genome replication in the presence of DelVG. (A) RNA from Fig. 4B was analyzed for *IFNB1* transcription by qPCR. (B) RNA was harvested from indicated viral stocks and subjected to a DelVG-specific qPCR for each indicated segment. Signal was normalized by a qPCR that measures total amount, deletion-containing and full-length, of each individual segment. Design of this qPCR was described in a prior publication.^41^ (C) A549 cells were infected, or not, with NS1_mut_ or a nominally wild-type population confirmed to have high DelVG content by qPCR at a genome-calibrated MOI of 0.4. Cells were treated, or not, with 50 μg/mL cycloheximide at the time of infection. RNA was harvested 8h post-infection and analyzed by qPCR for *IFNB1* transcripts normalized to *L32*. Controls demonstrating equivalent transcription under cycloheximide treatment in Fig S.23 (D) A549 cells were treated with 50 μg/mL cycloheximide at the time of infection and infected with nominally wild-type WSN confirmed to have high, or low, DelVG content at a genomecalibrated MOI of 1. Cells were additionally treated, or not, with 100 nM baloxaviric acid at the time of infection. RNA was harvested 8h post-infection and analyzed by qPCR. Measurements confirming silencing of *NELFB* and impacts on flu RNA levels in Fig S.24. (A-D) Significance assessed between silencing and NTC (A,D) strains (B) or cycloheximide treatment (C) by two-tailed Student’s t-test with Benjamini-Hochberg error correction, n=3, p<0.05. For all panels points represent biological replicates, lines represent mean values.

This finding was somewhat surprising, as prior reports indicate that under cycloheximide treatment viral transcription can contribute to interferon induction^77^. Two possibilities occurred to us: 1) viral transcription driven by NELF depletion and transcription that drives interferon induction are fundamentally different molecular events, or 2) there is some feature of flu populations that is largely absent from NS1_mut_ that determines interferon-stimulatory transcription. For the latter possibility, we wished to test whether transcription-dependent interferon production in turn depends on aberrant flu genomes, such as deletion-containing viral genomes (DelVGs). These are replication-incompetent genomes bearing large internal deletions that can accumulate in flu populations and are observed to correlate with an enhanced interferon response^10,14,15,17,41^.

To assess whether DelVG’s were absent from our NS1_mut_ stock, and therefore might explain this discrepancy, we used a DelVG-specific qPCR to measure their abundance in the three segments in which they are generally found (PB2, PB1, and PA)^41^. We find that indeed they are largely absent from this viral preparation, present at lower levels than even a stock we have previously characterized as “low” DelVG (Fig. 6B). When we compare infection with a viral stock prepared to have high DelVGs to NS1_mut_ virus, we find cycloheximide treatment enhances interferon production in response to infection with the former, but not the latter, consistent with prior results (Fig. 6C).^10,78^ We note that in general cycloheximide enhances interferon transcription through a cryptic mechanism.^79^ The observation with NS1_mut_ matches our own prior work and that of others that genome replication is a key step in interferon induction by this virus, but that DelVG viruses can induce a response in the absence of genome replication.^10,41^

Infecting cells during cyclohexamide treatment with viral stocks with high and low DelVG content, we find a strong role for viral transcription in triggering innate immunity only when DelVG content is high, and that silencing of *NELFB* enhances this effect (Fig. 6D). Curiously, we do still observe a slight enhancement of interferon production even upon baloxavir treatment, however the magnitude of effect is much smaller, and, unlike in the absence of baloxavir, there is little difference between high– and low-DelVG populations. We also see incomplete suppression of NELF-B-dependent increases in flu RNA in this experiment, perhaps due to incomplete blocking by baloxavir (Fig S.24). The very small increase (in absolute magnitude) in interferon transcription in our low DelVG population may be due to the slightly increased presence of deletions in the PB1 segment relative to our NS1_mut_ population (Fig 6B) Overall these data are consistent with the hypothesis that flu can generate transcription-dependent ligands, and that silencing of *NELFB* can increase their abundance. To demonstrate this is not an artifact of cycloheximide, nor of *NELFB* silencing, we confirm that infection with a characterized DelVG can induce interferon in a transcription-dependent fashion (Fig S.25). Therefore transcriptional increases upon NELFB depletion appears to enhance an interferon response both by supporting genome replication and by increasing the production of some transcription-dependent, DelVG-dependent, ligand.

## Discussion

Here we report the first transcription-based genome-wide CRISPR screen for genes that influence interferon induction. To accomplish this goal, we developed CRISPR with Transcriptional Readout, or CRITR-seq, in which each gRNA sequence serves as a barcode in a longer mRNA, associating the edit with transcription at a reporter promoter. No approach is perfect, and we should caveat the fact that we use a reporter at a non-endogenous locus – meaning we do not fully recapitulate all upstream enhancer elements nor post-transcriptional or post-translational controls of the targeted pathway. Nevertheless, despite this caveat we were still able to make novel findings that would not have been otherwise possible.

In applying CRITR-seq to identify host factors that modulate interferon induction by flu, we observed that many of the genes with the strongest effect sizes, both positive and negative, directly affect the flu life cycle. Gene edits that ablate interferon induction included not only members of the RIG-I signaling pathway, but also viral dependency factors required for host cell entry and the generation and nuclear export of *de novo* viral RNAs. Similarly, the top three hits for gene edits with enhanced interferon induction were all members of the NELF complex, which we have now defined as influencing flu transcription. These results suggest that some of the strongest observable host effects on interferon induction come from factors that impact the flu life cycle. This finding is consistent with our previous work which showed that in the absence of NS1, higher levels of interferon induction were associated with higher levels of flu RNA^10^. We do caveat that such effects are unlikely the sole drivers of the interferon response particularly given a recent preprint on OASL and interferon probability, which is also predicted to be a positive regulator of interferon in our screen here, as well as recent work describing host-derived ligands of RIG-I during flu infection.^80,81^

Our top hit, the NELF complex, has not previously been defined as a key host interface for influenza A virus. NELF depletion led to a massive increase in interferon induction at an early infection time point, which we found to be largely due to an increase in flu transcription and thus enhanced generation of immunostimulatory ligands. While loss of NELF accelerates transcription early in infection, it does not ultimately increase the total flu RNA levels or progeny virions. It seems that enhancement at this early step of RNA generation does not benefit the virus but instead increases opportunity for detection by cell intrinsic innate immunity, thereby disrupting mechanisms controlling progression through the viral life cycle that seem to minimize this detection.

It is surprising that NELF depletion increases viral RNA production even for a replication-incompetent virus or during cycloheximide treatment, as under these circumstances there are only 8 individual RNA molecules delivered by the initiating flu particle undergoing transcription at a given time. This suggests that establishing a productive interaction between the flu polymerase and the nascent capped host mRNAs is a rate-limiting process for viral RNA generation even when flu polymerase concentrations are incredibly low. Given that knockdown of the cap-binding complex also increases flu transcription, we speculate that the cap-binding complex, recruited by NELF^62,63,76^, competes with the flu polymerase for binding to nascent mRNA caps (Fig. 7A). Loss of NELF may expand the window in which capped mRNAs are available for flu polymerase binding by delaying both association of the cap-binding complex as well as the transition of RNA polymerase II to productive elongation^57^.

**Figure 7.**
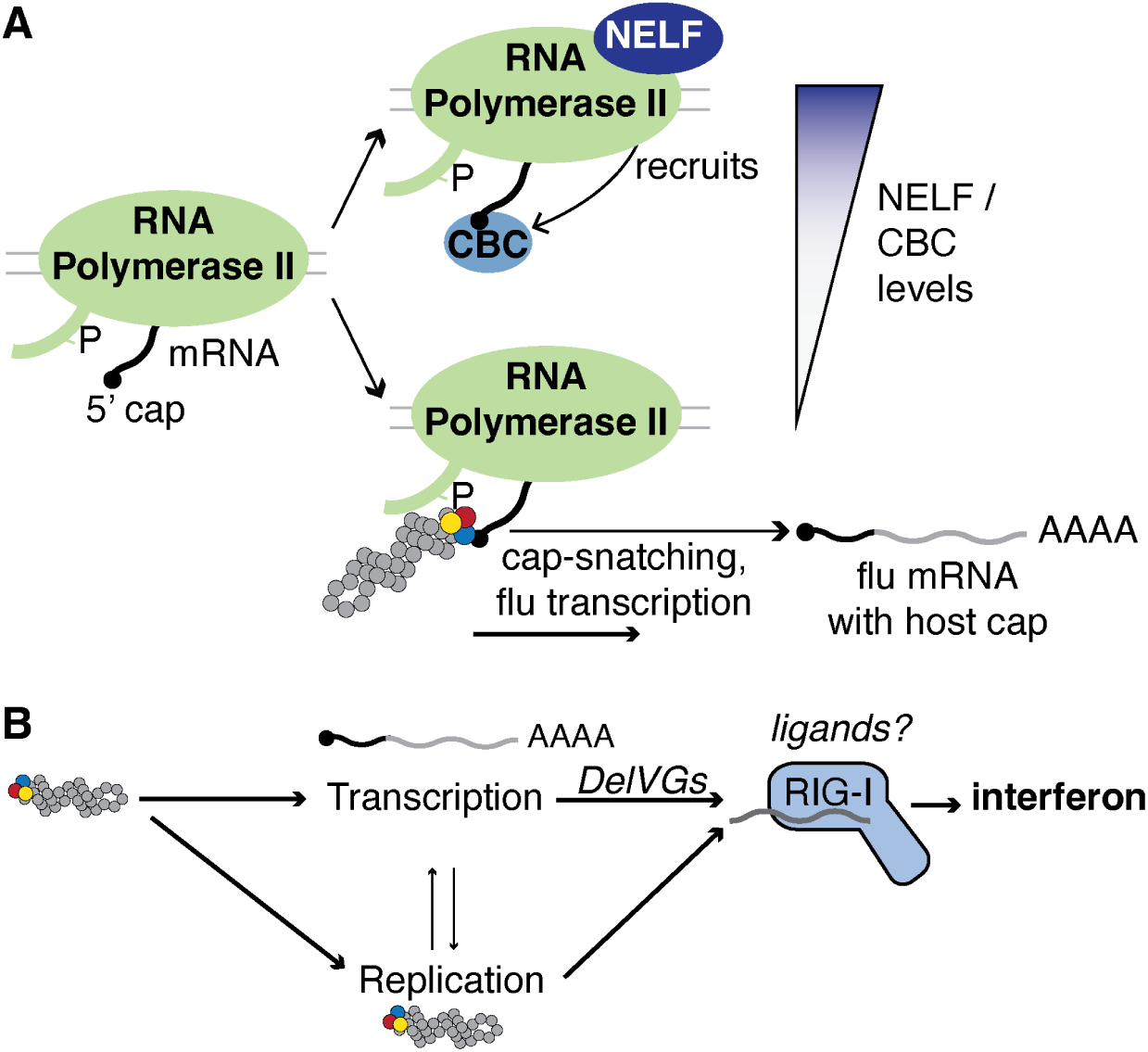
Model extrapolated from this study. (A) Influenza polymerase binds preferentially to RNA polymerase II with serine 5 phosphorylation in the C-terminal domain (CTD), which is the state that is characteristic of the promoter-proximal pause^61,64^. In our proposed mechanism, flu polymerase and cap-binding complex (CBC) compete for binding to nascent host mRNA caps. This is consistent with the finding in this study that depletion of NELF, which decreases CBC binding, increases flu cap-snatching and transcription. (B) Contributions of *de novo* viral RNA generation to interferon induction. Flu transcription contributes to interferon induction when DelVGs are present, although the transcription-dependent immunostimulatory ligand(s) remain unknown. Flu replication also contributes to interferon induction, and is directly influenced by rates of transcription (i.e., viral genomes).

Many previous studies have demonstrated the importance of *de novo* viral RNA for interferon induction by flu^10,16,41,77^. Our finding, in the context of NELF depletion, that increased transcription leads to an increase in interferon induction demonstrates that flu transcription is not only critical but also rate-limiting for the interferon response. This appears to include interferon induction when viral replication is blocked. Although genome replication products have been generally regarded as the primary influenza ligand, our finding is consistent with previous work from Richard Randall’s group, which also demonstrated transcription-dependent, replication-independent interferon induction^77^. Thus, we conclude that influenza transcription-dependent processes, independent of genome replication, contribute to interferon induction (Fig. 7B), although our work extends that conclusion in that DelVGs, or other aberrant viral genomes, appear to be critical for generation of such ligands.

Determining which flu RNA(s) are the relevant immunostimulatory ligands in flu infection remains to be clarified, although it seems increasingly likely that multiple ligands, both host– and virally-derived, exist. Both transcription and replication have now been conclusively shown to be critical steps for interferon induction. Here, we show that transcription itself can contribute to interferon induction, at least in the case of deletion-containing defective genomes, which can be abundant even in “wild-type” flu populations^82,83^. This may explain why some defective viral particles induce interferon at such high levels, despite only having 8 viral genomes in the entire cell. However, it is unclear what the transcription-dependent, replication-independent ligand(s) may be, as it seems unlikely that flu mRNAs, which are capped and polyadenylated like host mRNAs, would activate RIG-I. Recent work by the te Velthuis lab demonstrated that the flu polymerase can generate capped cRNAs (ccRNAs), which in complex with other small viral RNAs contribute to the interferon response^84^. Generation of ccRNAs would likely require cap-snatching and may be the transcription-dependent ligand we observe here.

In conclusion, a CRITR-seq screen for regulators of interferon expression led us to the finding that components of the host transcription machinery – NELF and the cap-binding complex – modulate the rate of flu transcription and thus the interferon response. In the future, we hope the CRITR-seq platform will be useful to other studies of phenotypes related to gene expression, simply by swapping out the interferon promoter we used here for another promoter of interest.

## Materials and Methods

### Cells

The following cell lines were used in this study: HEK293T (human embryonic kidney, female; ATCC CRL-3216), MDCK-SIAT1 (variant of the Madin Darby canine kidney cell line overexpressing SIAT1, female cocker spaniel; Sigma-Aldrich 05071502), Huh-7 (human hepatocellular carcinoma; JCRB0403)and A549 (human lung epithelial carcinoma, male; ATCC CCL-185).

Cells were cultured in D10 media (Dulbecco’s Modified Eagle Medium supplemented with 10% heat-inactivated fetal bovine serum and 2 mM L-Glutamine) in a 37°C incubator with 5% CO_2_. Cell lines were tested for mycoplasma using the LookOut Mycoplasma PCR Detection Kit (Sigma-Aldrich) with JumpStart Taq DNA Polymerase (Sigma-Aldrich), or using the MycoStrip Mycoplasma Detection Kit (InvivoGen, rep-mys-20).

Generation of the specific A549 cell lines (type III interferon reporter, A549-Cas9-mCherry, and *RIG-I* –knockout lines) used in this study are described in supplementary information.

### Viruses

Genomic sequences for wild-type A/WSN/1933 (H1N1), NS1_stop_, NS1_mut_, and PB1_455:350_ influenza viruses are in File S2. Flu viruses were generated by reverse genetics^85^ and titered by TCID_50_ on MDCK-SIAT1 cells or qPCR against the viral segment HA. For wild-type WSN, the virus generated by reverse genetics was then expanded by infecting MDCK-SIAT1 cells at an infectious MOI of 0.01, in order to obtain a population with a low abundance of deletion-containing genomes. Detailed methods, including coronavirus OC43 propagation, are described in supplementary information.

### Genome-Wide CRITR-seq Screen

Detailed protocol for the CRITR-seq screen is described in supplementary information. In brief, Guides from the Human Genome-Scale CRISPR Knockout (GeCKO) v2 Pooled Library (Addgene Pooled Libraries #1000000048, #1000000049)^39^ were cloned into the CRITR-seq vector. CRITR-seq plasmid libraries were transfected into HEK293T cells to generate lentiviral libraries. A clonal Cas9-expressing A549 line was transduced with the lentiviral libraries at an MOI of 1.5. This is a higher MOI than is typical for CRISPR screens, as we assumed that most gene edits would not impact interferon expression. On day 10 post-transduction, cells were infected with NS1_mut_ at a genome-corrected MOI of 2. 8 hours post-infection, cells were lysed for RNA and genomic DNA extraction. The region containing the gRNA sequence in the CRITR-seq vector was PCR-amplified from all samples and prepared for amplicon sequencing. The entire process from PCR to final gDNA/RNA extraction and sequencing preparation was repeated 3 times for 3 independent biological replicates of the CRITR-seq screen.

After amplicon sequencing, Model-based Analysis of Genome-wide CRISPR/Cas9 Knockout (MAGeCK)^38^ was performed to identify genes with gRNAs enriched or depleted in the mRNA relative to the gDNA. Parameters for sequencing analysis are described in supplementary information.

### Individual Gene Knockout by CRISPR/Cas9

Individual gene knockouts were performed by CRISPR ribonucleoprotein (crRNP) transfection, using the Alt-R S.p. HiFi Cas9 Nuclease V3 (Integrated DNA Technologies, 1081060). Alt-R CRISPR-Cas9 tracrRNA (Integrated DNA Technologies, 1073189) and the specific crRNA were complexed together by mixing at equimolar concentration to a final concentration of 1 μM in Nuclease-Free Duplex Buffer (Integrated DNA Technologies, 0000934164) and heating at 95°C for 5 minutes. The RNP complex was formed by mixing the crRNA-tracrRNA duplex and Cas9 to a final concentration of 60 nM each in Opti-MEM and incubating 5 minutes at room temperature. The crRNP complex was reverse transfected into 160,000 A549 cells in a 24-well plate using Lipofectamine RNAiMAX (Invitrogen, 100014472), with a final concentration of 6 nM crRNP. Experiments were performed 10 days post-transfection to allow sufficient time for editing and protein turnover. All gene-specific guides used for these knockouts were independent of guides present in the GeCKO library (TableS4).

### siRNA Knockdown

All siRNAs were Ambion Silencer Select siRNAs (Life Technologies Corporation). Catalog numbers are provided in Table S4. For *NELFB* knockdown, 100,000 A549 cells/well in a 24-well plate were each reverse-transfected in 500 μL D10 with 50 μL Opti-MEM, 1.5 μL of Lipofectamine 3000 (Invitrogen, L3000001), and 0.25 μL of 10 μM siRNA. Media was replaced with fresh D10 24 hours post-transfection. 4 days after transfection, cells were trypsinized and again reverse-transfected with siRNA. siRNA treatment was repeated again 4 days after the second transfection to seed for infection at 100,000 cells/well. Influenza infection or poly(I:C) transfection was performed 24 hours after the third transfection. *NCBP1* knockdown was performed based on the methods from Gebhardt et al.^86^, as described in supplementary information. *CSNKB2B* silencing further described in supplementary information.

### Poly(I:C) Transfection

500 ng poly(I:C) (Fisher Scientific, 42-871-0) was complexed with 1.5 μL Lipofectamine 3000 in 50 μL Opti-MEM. 5 μL of the complex (containing 50 ng poly(I:C)) was added to each well of a 24-well plate, with 100,000 A549 or 293T cells/well. Cells were lysed for RNA extraction 8 hours post-transfection.

### qPCR

Infections were performed on cells seeded 24 hours prior to infection, changed to Influenza Growth Medium (IGM, Opti-MEM supplemented with 0.04% bovine serum albumin fraction V, 100 μg/ml of CaCl2, and 0.01% heat-inactivated fetal bovine serum) at time of infection. RNA was purified from infected cells or viral supernatant using the Monarch Total RNA Miniprep kit from New England Biolabs or the Zymo Research Quick-RNA Miniprep kit, following manufacturer’s protocol. cDNA from 100 ng purified RNA was generated using the High Capacity cDNA Reverse Transcription Kit (Applied Biosystems, 4368814) with random hexamer primers, oligo(dT) primers (for initial screen validation, Fig. 3B), or universal influenza primers (for viral titer from supernatant, Fig. 4E, S20) according to the manufacturer’s protocol. When 100 ng purified RNA was not available (in viral titer experiments), we used a final lysate concentration of 10% of the reaction volume. qPCR was performed using Luna Universal qPCR Master Mix (New England Biolabs, M3003) with manufacturer’s suggested reaction conditions with no modifications. Primers for all qPCR analyses are listed in Table S5. Code generating graphs in manuscript can be found at https://github.com/Russell-laboratory/CRITRseq_Interferon_Flu. Transcripts encoding the human ribosomal protein L32, *L32*, were used as a housekeeping control, we have consistently observed similar CT values for this transcript even during flu infection, which otherwise can globally depress the host transcriptome.

### Inhibitor Treatment

Inhibitors were diluted in IGM to final concentrations of 10 nM or 100 nM for baloxaviric acid (Fisher Scientific 501872233), 100 nM for pimodivir (Fisher Scientific 502028875), or 50 μg/mL for cycloheximide (Fisher Scientific NC1163264). At time of infection, or 2 hours pre-infection for pimodivir experiments, 100,000 A549 cells in a 24-well plate were treated by replacing D10 from seeding with 500 μL IGM with or without the indicated inhibitors.

### Flow Cytometry

Indicated cells were seeded 24 hours prior to infection, changed to IGM at the time of infection, and trypsinized and resuspended in 1x phosphate-buffered saline (PBS) supplemented with 2% of heat-inactivated fetal bovine serum (FBS) 13 hours post-infection. For M2 staining, a mouse monoclonal antibody (Invitrogen, MA1-082) was used at a concentration of 5 μg/mL. Secondary antibody staining was performed with goat anti-mouse IgG conjugated to allophycocyanin (Invitrogen, A-865) at a concentration of 5 μg/mL. Cells were fixed with BD Cytofix/Cytoperm (BD Biosciences, 51-2090KZ), following manufacturers instructions.

FlowJo was used to generate a gate excluding debris prior to export to a csv. Data for non-debris events were analyzed using custom Python scripts, which can be found at https://github.com/Russell-laboratory/CRITRseq_Interferon_Flu. Uninfected controls were used to set empirical gates for influenza staining and interferon reporter positivity at a 99.9% exclusion criteria.

### 5’ Rapid Amplification of cDNA Ends (RACE)

To identify the sequences of the 5’ terminal ends of viral mRNA, samples were reverse transcribed with the Template Switching RT Enzyme Mix (New England Biolabs, #M0466) according to the manufacturer’s protocol. Amplification of the 5’ terminal ends of the flu mRNA were performed in separate reactions using a template switch oligo (TSO)-specific amplification primer and segment-specific mRNA primers (Table S5)^89^. Detailed methods for 5’ RACE sample preparation and sequencing analysis are described in supplementary information.

### Single-cycle Viral Titer

A549 cells were seeded at 100,000 cells/well in a 24 well plate. 24 hours after seeding, the media was replaced with IGM and infected with WT WSN at an infectious MOI of 2. 2 hours post-infection, the media was replaced with fresh IGM. 14 hours post-infection, supernatant was collected and centrifuged at 300g for 4 minutes to clear cell debris. 40 μL of supernatant was mixed with 360 μL RNA lysis buffer for qPCR analysis. Cells were also lysed for RNA extraction and qPCR analysis.

### Western Blot

After 24 hour infection by NS1_mut_ at a genome-corrected MOI of 5, cells were lysed, lysate was resolved by SDS-PAGE, and RIG-I and β-actin were probed by Western blot. For NELF-B measurements, cells were infected and treated as in Fig. 3C for 8h, and cells were lysed, lysate was resolved by SDS-PAGE, and NELF-B and β-actin were probed by Western blot. Detailed methods are described in supplementary information.

## Materials and Data Availability

Sequencing data have been deposited at Gene Expression Ominbus (GEO) as GEO: GSE281730 and are publicly available. All original code, as well as qPCR data, have been deposited at https://github.com/Russell-laboratory/CRITRseq_Interferon_Flu and are publicly available. Plasmids and cell lines generated for this study are available from the corresponding author, Alistair B. Russell.

## Acknowledgments

This work was supported by the NIGMS of the NIH under grant R35GM147031 awarded to ABR. ACV was supported by the NIGMS of the NIH under grant T32GM133351. Work performed by SNZJ was supported in part by the NIAID of the NIH under grant 5R01AI176639 awarded to ERT. We thank Scott Biering and Matt Daugherty for help and support with work on coronavirus OC43, and Emily Troemel for advice and support in preparing this work.

## Author contributions

A.B.R. and A.C.V. designed experiments, A.B.R., A.C.V, S.N.Z.J., M.M., S.S., L.K.C., J.S.P, and K.D.V. performed experiments, A.B.R and A.C.V performed data analysis, A.B.R and A.C.V. conceived this study, discussed results, and wrote the manuscript, and S.N.Z.J, M.M., S.S., and K.D.V. provided thoughtful feedback on drafts of the manuscript.

## Supplementary Information

### Supplementary Methods

#### Generation of A549 cell lines

The parental A549 cell line used to make the type III interferon reporter line, A49-Cas9-mCherry line, and *RIG-I* – knockout line was authenticated using the ATCC STR profiling service.

A549 type III interferon reporter line was previously described^40^.

A clonal A549 cell line constitutively expressing Cas9 (A549-Cas9-mCherry) was generated by transducing A549 cells with a lentiviral vector containing Cas9 under the control of a CMV promoter (plasmid sequence in File S1). The mCherry sequence is connected to the Cas9 mRNA by a P2A linker. Cells were validated to be free of replication-competent lentivirus via qPCR for VSV-G, primers in Table S5. Single cells containing the Cas9 expression construct were isolated by fluorescence-activated cell sorting for mCherry-positive cells. The final clonal line was validated by flow cytometry to have unimodal, high levels of mCherry expression.

An A549 *RIG-I* –knockout cell line was generated by transfecting A549 cells with a Cas9 ribonucleoprotein complex, using a CRISPR RNA (crRNA) targeting *RIG-I* exon 1 (IDT Alt-R CRISPR-Cas9 crRNA Hs.Cas9.DDX58.1.AA, Table S4). Single cells were isolated by dilution cloning to generate a clonal line. Residual RIG-I protein expression in this cell line is shown in Figure S21.

#### Viruses

Wild-type A/WSN/1933 (H1N1) influenza virus was created by reverse genetics using plasmids pHW181-PB2, pHW182-PB1, pHW183-PA, pHW184-HA, pHW185-NP, pHW186-NA, pHW187-M, pHW188-NS^85^. Genomic sequence of this virus is provided in File S2. HEK293T and MDCK-SIAT1 cells were seeded in an 8:1 coculture and transfected using BioT (Bioland Scientific, LLC) 24 hours later with equimolar reverse genetics plasmids. 24 hours post-transfection, D10 media was changed to Influenza Growth Medium (IGM, Opti-MEM supplemented with 0.04% bovine serum albumin fraction V, 100 μg/mL of CaCl_2_, and 0.01% heat-inactivated fetal bovine serum). 24 hours after the media change, viral supernatant was collected, centrifuged at 300g for 4 minutes to remove cellular debris, and aliquoted into cryovials to be stored at –80°C. Thawed aliquots were titered by TCID_50_ on MDCK-SIAT1 cells, and calculated using the Reed and Muench formula^87^. To generate a viral population with a low abundance of deletion-containing genomes, the virus generated by reverse genetics was then expanded by infecting 3×10^6^ MDCK-SIAT1 cells in a 10 cm plate at an infectious MOI of 0.01. Viral supernatant was harvested at 36 hours post-infection, stored, and titered in the same way as the original virus.

NS1_stop_ and NS1_mut_ viruses were generated by reverse genetics using the same plasmids for genome segments 1-7 of the A/WSN/1933 WT virus, but replacing pHW188-NS with a variant NS sequence of pHW188-NS (NS1_stop_) encoding the NS segment from WSN, or of pHW198 (NS1_mut_), encoding the NS segment of PR8. These variant sequences have 3 stop codons early in the open reading frame of NS1, and the sequences are included in File S2. HEK293T cells were cocultured with MDCK-SIAT1 cells constitutively expressing NS1 from PR8, and the coculture was transfected with the 8 reverse genetics plasmids as well as a pHAGE2 plasmid containing the gene for WSN-NS1 under the control of the constitutive CMV promoter. D10 was changed to IGM 24 hours post-transfection, and virus was harvested 72 (NS1_stop_) or 52 (NS1_mut_) hours post-transfection. Thawed aliquots were titered by TCID_50_ on MDCK-SIAT1 cells constitutively expressing NS1, or by qPCR against the viral segment HA.

To generate PB1_455:350_^41^, HEK293T cells were cocultured with MDCK-SIAT1 cells constitutively expressing PB1. The coculture was transfected with all reverse-genetics plasmids encoding A/WSN/1933 except for a variant of pHW182-PB1 with the indicated deletion (File S2), and a construct expressing PB1 under control of a constitutive promoter in the pHAGE2 vector. D10 was changed to IGM 24 hours post-transfection, and virus was harvested 72 hours after the media change. This virus was titered on MDCK-SIAT1 cells constitutively expressing PB1.

To generate a high-DelVG wild-type virus population^10^, reverse-genetics was performed with the 8 WSN plasmids in every well of a 96-well plate. After 48h, 10 μL of supernatant was transferred to fresh wells, each seeded with 10,000 MDCK-SIAT1 cells. After 48h, 10 μL of that supernatant was transferred to fresh wells, each seeded with 10,000 MDCK-SIAT1 cells. After 48h, all supernatants were harvested, pooled, and clarified, to generate one biological replicate viral population. This virus was titered by TCID_50_ on MDCK-SIAT1 cells, and by qPCR against the viral segment HA.

The original stock of coronavirus OC43 used in this work was provided by Scott Biering. This stock was amplified to produce virus used in this work by infecting Huh-7 cells at 70% confluence in D10 media at 32°C for two days. Cells and supernatant were then harvested, using a cell scraper to resuspend adherent cells. Supernatant was clarified by centrifugation at 300g for 4 minutes and set aside. Cells were subjected to two rounds of freeze-thaw at –80°C. Superanatant and cells were recombined, and resultant mixture re-clarified by centrifugation at 300g for 4 minutes, aliquoted, and stored at –80°C prior to use. OC43 was titered on A549 cells by infection at 32°C with various dilutions, and staining with OC43 nucleocapsid antibody (Cat, 40643-T62, SinoBiological) at 24h post-infection.

#### Genome-wide CRITR-seq screen

Guides from the Human Genome-Scale CRISPR Knockout (GeCKO) v2 Pooled Library^39^ (Addgene Pooled Libraries #1000000048, #1000000049) were cloned into the CRITR-seq vector in triplicate reactions. To minimize jackpotting from PCR, gRNAs were amplified from GeCKO half-libraries A and B in 8 (Library A) or 7 (Library B) separate 20-cycle PCR reactions, pooled by library, and purified by gel extraction. See Table S5 for primers used in PCR reactions. gRNAs were inserted into the CRITR-seq vector backbone (plasmid sequence in File S3) by Gibson assembly using NEBuilder HiFi DNA Assembly Master Mix (New England Biolabs, E2621), with a 15:1 insert:backbone molar ratio and a 1 hour incubation. DNA >∼1000 bp was purified from the reaction mix using ProNex Size-Selective Purification System magnetic beads (Promega NG2002) at 1x bead volumes, and electroporated into ElectroMAX^TM^ Stbl4^TM^ Competent Cells (Invitrogen 11635018) to achieve at least 10x as many colonies as guides for each half-library. Colonies were scraped from plates, incubated in 100 mL LB at 30°C for 1 hour with shaking, and plasmids were purified by midiprep. When needed, CRITR-seq plasmid libraries were amplified by electroporating 4 ng of the original generated library.

To generate lentiviral libraries, 6.32×10^6^ HEK293T cells were seeded in D10 in each of 2 T75 flasks to achieve ∼80% confluence. Cells in each flask were transfected with a total of 10.27 μg DNA combined from CRITR-seq libraries A and B, proportional to the number of guides per library. Plasmid libraries were transfected using Lipofectamine 3000 and P3000, along with lentiviral helper plasmids HDM-Hgpm2 (Addgene #204152), HDM-tat1b (Addgene #204154), pRC-CMV-Rev1 (Addgene #164443), and HDM_VSV_G (Addgene #204156), at 10.27 μg divided equally between them. 6 hours post-transfection, the media was changed to fresh D10. 24 hours post-transfection, virus was harvested and stored at 4°C, replacing the media. 50 hours post-transfection, virus was harvested again, pooled with the first harvest, and centrifuged at 300 g for 4 minutes to remove cell debris. Aliquots were stored at –80°C. The CRITR-seq vector does not contain a selectable marker, so we titered the lentiviral libraries by flow cytometry, taking advantage of the zsGreen marker under the control of the *IFNL1* promoter. A549 cells were seeded at 67,500 cells/well in a 24-well plate and transduced with serial dilutions of lentivirus from 1x to 256x, with 3 replicates per dilution. 24 hours post-transduction, the media was replaced with fresh D10. 48 hours post-transduction, cells were seeded for infection at 70,000 cells/well. Cells were infected at a saturating infectious MOI of 2.8 using an influenza variant that induces interferon in ∼100% of infected cells, and fixed with formaldehyde 13 hours post-infection. The percent of zsGreen+ cells measured by flow cytometry was considered to be the fraction of cells that had integrated the CRITR-seq vector. We used the dilutions which appeared to be in the linear range to calculate the transduction units (TU) per μL for each library.

To generate libraries of edited cells, 2.666×10^6^ A549-Cas9-mCherry cells were seeded in each of 3 T75 flasks. Cells were transduced with a CRITR-seq lentiviral library at an MOI of 1.5, using 8 μg/mL polybrene (Sigma-Aldrich, TR-1003-G). This is a higher MOI than is typical for CRISPR screens, as we assumed that most gene edits would not impact interferon expression. With this MOI and cell count, we were transducing cells with ∼100x coverage of the gRNAs from the GeCKO library. 24 hours post-transduction, the media was changed to fresh D10, and cells were passaged for 10 days, maintaining over 100x coverage of the libraries at each passage. On day 10 post-transduction, cells were seeded at 7×10^6^ cells per 15 cm plate in 12 plates for ∼1000x coverage of gRNAs. The day following seeding, D10 was replaced with IGM for infection with NS1_mut_ at a genome-corrected MOI of 2. 8 hours post-infection, cells from all plates were trypsinized and pooled. Half of the cells were lysed in genomic DNA lysis buffer and processed for genomic DNA extraction (Monarch Spin gDNA Extraction Kit; New England Biolabs, T3010S), and half of the cells were lysed in RNA lysis buffer and processed for RNA extraction (Monarch Total RNA Miniprep Kit; New England Biolabs, T2010S).

To prepare genomic DNA for amplicon sequencing, 50 μg gDNA was split into 20 50-μL PCR reactions. Assuming 6 pg gDNA per cell and 1.5 guides per cell, this would give ∼100x coverage of gRNAs. We used Q5 Hot Start High-Fidelity 2X Master Mix and primers specific for a 194 base pair region of the CRITR-seq vector containing the gRNA sequence (Table S5). Amplicons were bead purified using 3x bead volumes, and 5% of the bead-cleaned product was amplified by a second PCR using primers containing partial Illumina adapters (Table S5). Amplicons were purified in a 2-sided bead clean-up, and DNA concentrations were determined by Qubit. Final Illumina indices were added by PCR for each sample, using 10 ng of amplicon. Indexing primers were from IDT for Illumina DNA/RNA unique dual indexes (Set A, #20026121). PCR reactions were bead purified using 2x bead volumes and verified by running on an agarose gel. Final DNA concentrations were determined by Qubit before pooling equal amounts for sequencing. Our initial, low-depth, sequencing was performed as follows; we reverse transcribed 1 μg of total RNA using oligo(dT) primers and the Superscript III First-Strand Synthesis SuperMix kit (Invitrogen, 18080400) according to the manufacturer’s protocol. 10% of the total RT reaction was amplified in each of 5 PCR reactions. PCR amplification and indexing was performed with the same steps and primers as used for genomic DNA preparation. Subsequent re-analysis of data, including a molecularly-calibrated qPCR using primers to generate the first PCR product in our process revealed that our input into our initial sequencing was likely only ∼120,000 molecules, effectively sampling the library at 1x coverage. This qPCR used our sequencing primers, under identical PCR conditions, and a plasmid control, to measure the precise number of molecules that went into our sequencing run. We repeated cDNA generation using a gene-specific primer specified in Table S5, which we found to improve recovery of cDNA ∼two-fold with 5 μg of total RNA in a scaled-up reaction with Superscript III First-Strand Synthesis SuperMix kit, following the manufacturer’s protcol. The same molecularly-calibrated qPCR inferred that this would result in 40-fold coverage of our libraries, at an estimated 4,800,000 molecules of input. The entirety of this reaction was then amplified using Q5 polymerase and comprising no more than 10% of the PCR volume using primers specified in Table S5 with the following reaction conditions: 59°C annealing, 15s extension, and 28 cycles. Reactions were gel-extracted, and 10ng of the resultant product subjected to an additional 8 rounds of indexing using Illumina index primers. Data from this are presented throughout the manuscript, initial low-quality sequencing was only used to generate data used to select for our secondary screen. Additional instances of CRITR-seq should use this protocol.

The entire process from PCR to final gDNA/RNA extraction and sequencing preparation was repeated 3 times for 3 independent biological replicates of the CRITR-seq screen.

For our secondary screen, we ordered guides synthesized from Twist Biosciences as described in Table S7 and cloned into our CRITR-seq vector three independent times to generate our three replicates. We performed CRITR-seq as in our primary screen with the following modifications: only 6×10^6^ 293tT cells were used per library to generate lentivirus with plasmids scaled appropriately, an MOI of 0.4 rather than 1.5 was used to reduce conflict, 3×10^6^ A549 cells were transduced in a 10cm plate, and for infection 28×10^6^ A549 cells were infected with our mutant flu virus.

#### siRNA knockdown of *NCBP1* and *CSNK2B*

siRNA sequence and protocol for *NCBP1* knockdown were based on the methods from Gebhardt et al.^86^. 100,000 A549 cells/well in a 6-well plate were each reverse-transfected in 2 mL D10 with 250 μL Opti-MEM, 7.5 μL of Lipofectamine RNAiMAX, and 2.5 μL of 10 μM siRNA. Media was replaced with fresh D10 24 hours post-transfection. 48 hours after transfection, cells were trypsinized and re-treated with siRNA, at 200,000 cells/well in a 6-well plate. 24 hours after the second transfection, cells were trypsinized and re-seeded in a 24-well plate at 50,000 cells/well, with no additional siRNA treatment. Influenza infection was performed 24 hours after seeding (96 hours after the first transfection).

Silencing of *CSNK2B* was performed near-identically, however silencing was only performed for 24h, at a single dose, prior to infection, rather than the longer silencing approach used for *NCBP1*.

#### Complementation of *NELFB*

Lentivirus expressing codon-optimized *NELF-B* was generated by seeding 4×10^5^ HEK293T cells in a 6-well plate with D10, followed by transfection with 1 μg DNA from a plasmid encoding codon-optimized *NELFB* (sequence provided in File S4), and 1 μg DNA consisting of equal amounts of the lentiviral helper plasmids HDM-Hgpm2 (Addgene #204152), HDM-tat1b (Addgene #204154), pRC-CMV-Rev1 (Addgene #164443), and HDM_VSV_G (Addgene #204156). 24h post-transfection, media was changed to fresh D10. 72h post-transfection, supernatant was harvested, syringe-filtered, and used for transduction.

A549 cells were treated with silencing RNA targeting *NELFB* or a non-targeting control as in other qPCR experiments. On day 8 of silencing, cells were not only re-treated with siRNA, as in other experiments, but some were transduced with our *NELFB* expression vector by adding lentiviral supernatant and polybrene to a final concentration of 5μg/ml. 19.5h post-transduction media was changed to IGM and cells were infected and analyzed as described in Fig S.11.

To generate lentiviral libraries, 6.32×10^6^ HEK293T cells were seeded in D10 in each of 2 T75 flasks to achieve ∼80% confluence. Cells in each flask were transfected with a total of 10.27 μg DNA combined from CRITR-seq libraries A and B, proportional to the number of guides per library. Plasmid libraries were transfected using Lipofectamine 3000 and P3000, along with lentiviral helper plasmids HDM-Hgpm2 (Addgene #204152), HDM-tat1b (Addgene #204154), pRC-CMV-Rev1 (Addgene #164443), and HDM_VSV_G (Addgene #204156), at 10.27 μg divided equally between them. 6 hours post-transfection, the media was changed to fresh D10. 24 hours posttransfection, virus was harvested and stored at 4°C, replacing the media with fresh D10. 50 hours post-transfection, virus was harvested again, pooled with the first harvest, and centrifuged at 300 g for 4 minutes to remove cell debris.

#### 5’ Rapid Amplification of cDNA Ends (RACE)

To identify the sequences of the 5’ terminal ends of the viral mRNA upon NELFB knockdown, cells were treated with either no, non-targeting, or one of two *NELFB* siRNAs for 9 days according to the *NELFB* siRNA knockdown procedure described above. Cells were infected with NS1_mut_ at a genome-corrected MOI of 2 for 8 hours before lysing cells and extracting RNA using the Monarch Total RNA Miniprep Kit from New England Biolabs. Samples were reverse transcribed with the Template Switching RT Enzyme Mix (New England Biolabs, #M0466) according to the manufacturer’s protocol. In brief, sample RNA was annealed with oligo(dT) primers and then reverse transcribed with the template-switching oligo (TSO, Table S5), which contains mixed ribonucleotides at the third from final position, a locked nucleic acid at the final position, and a 5’ biotin modification to prevent spurious additional concatemerization, similar to that from Picelli *et al*.^88,89^. Amplification of the 5’ terminal ends of the flu mRNA were performed in separate reactions using a TSO-specific amplification primer and segment-specific mRNA primers (Table S5), using Q5 Hot Start High-Fidelity 2X Master Mix. Amplification was repeated with primers containing partial Illumina i5 and i7 adapters (common TSO partial adapter and segment-specific primers in Table S5). Amplicons were bead purified, pooled by sample, and verified on a gel. DNA concentration was determined by Qubit, and final Illumina indices were added by PCR for each sample, using 1.5 ng of amplicon. Indexing primers were from IDT for Illumina DNA/RNA unique dual indexes (Set A, #20026121). PCR reactions were bead purified, and final DNA concentrations were determined by Qubit before pooling equal amounts for sequencing on the Illumina Novaseq platform.

Reads containing a perfect match to the TSO were searched for the first 20 bases of the mRNA sequence from each of the 8 flu genome segments. Reads with at least 15 nucleotide identity were considered matching to that flu mRNA. The sequence downstream of the TSO and upstream of the +1 position of the viral mRNA sequence was considered to be the cap sequence. We note that the length of this sequence could be +/-1 compared to the true length of the cap sequence due to the variable number of non-templated nucleotides added by the reverse transcriptase after it reaches the 5’ end of the RNA. A large number of duplicate cap sequences indicated that there was substantial bottlenecking in the sample preparation process, so only unique cap sequences were kept for analysis. Median cap sequence length for each sample was 12 nucleotides. Distribution of cap sequence lengths were plotted, excluding the top 2% of lengths. Distributions were compared by one-way ANOVA and post hoc Tukey’s test, n=8, p<0.05, q<0.05.

#### Western blot

For RIG-I Western blot cells were seeded in a 24-well plate at 100,000 cells/well 24 hours prior to infection. Cells were infected with NS1_mut_ at a genome-corrected MOI of 5 for 24 hours. For NELF-B western blot, cells were seeded and infected identically to qPCR experiments. Infected cells were then washed three times with 1x PBS and lysed in 1x Laemmli Sample Buffer (Bio-Rad #1610747) containing 100 mM DTT. Lysate was boiled at 100°C for 5 minutes and resolved by SDS-PAGE (Bio-Rad #4561024) and transferred onto a nitrocellulose membrane. The membrane was blocked with PBST buffer (1x PBS with 0.1% Tween-20) containing 5% nonfat dry milk for 1 hour at room temperature. The membrane was incubated overnight at 4°C with anti-RIG-I monoclonal antibody (Cell Signaling Technology, 3743T; 1:1000 dilution in PBST), anti-NELF-B (Cell Signaling Technology, 14894T, 1:1000 dilution in PBST) or anti-β-actin monoclonal antibody (Cell Signaling Technology, 8457T; 1:1000 dilution in PBST) as a loading control. The membrane was then incubated for 1 hour at room temperature with HRP-linked anti-rabbit IgG secondary antibody (Cell Signaling Technology, 7074P2; 1:3000 dilution in PBST) to detect primary antibody staining. HRP-conjugates were visualized using the Clarity Western ECL Substrate kit (Bio-Rad #1705060S).

#### Viability assay

Viability was assessed using the CellTiter 96 AQ_ueous_ One Solution Cell Proliferation Assay (Promega # G3580) using manufacturer’s instructions. A549 cells were treated with siRNA against *NELF-B* or a non-targeting control as for other assays described in methods. On day 8 of silencing, when cells were re-treated with siRNA and reseeded, cells were seeded in a 96-well plate at a density of 25,000 cells/well in 100 μl of D10. To measure viability, on day 9 of silencing wells containing silenced cells, or an equivalent number of media controls, were treated by adding 20 μl of assay reagent, which contains 3-(4,5-dimethylthiazol-2-yl)-5-(3-carboxymethoxyphenyl)-2-(4-sulfophenyl)-2H-tetrazolium, or MTS. After 4h of incubation at 37°C with 5% CO_2_, the reduction of MTS to a formazan product was measured by platereader using absorbance at 490 nm.

MTS can be reduced by cells to a formazan product, which we measured at 490 nm on a plate-reader at 4h of incubation under standard tissue culture conditions (5% CO_2_). From this signal, we subtracted absorbance at 700 nm, indicative of nonspecific signal in our assay, to provide the final data presented in Fig. 3D.

## Supplementary Material

**Supplementary Figure 1.**
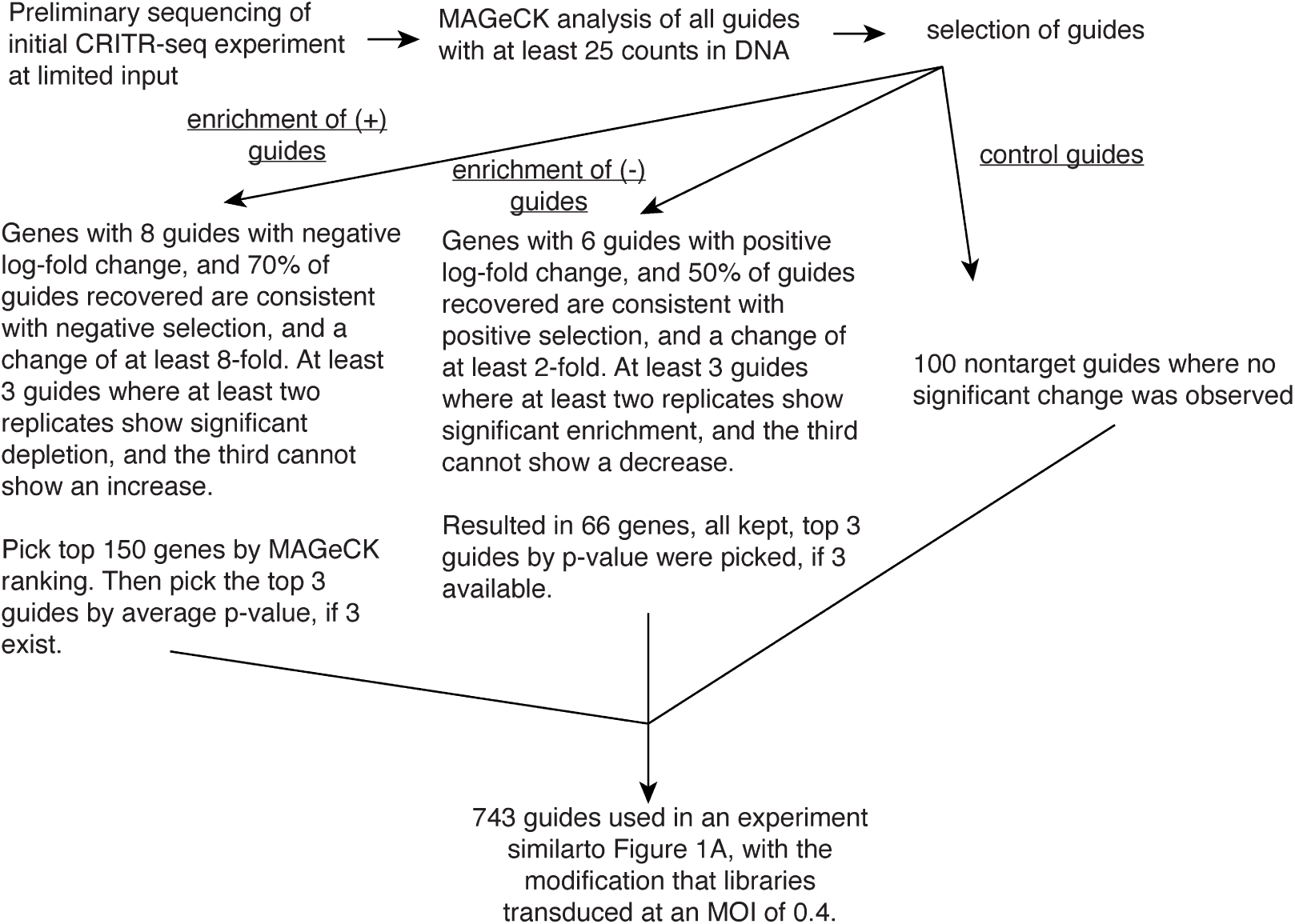
Workflow describing the smaller, secondary, screen used to assess replicability of CRITR-seq method.

**Supplementary Figure 2.**
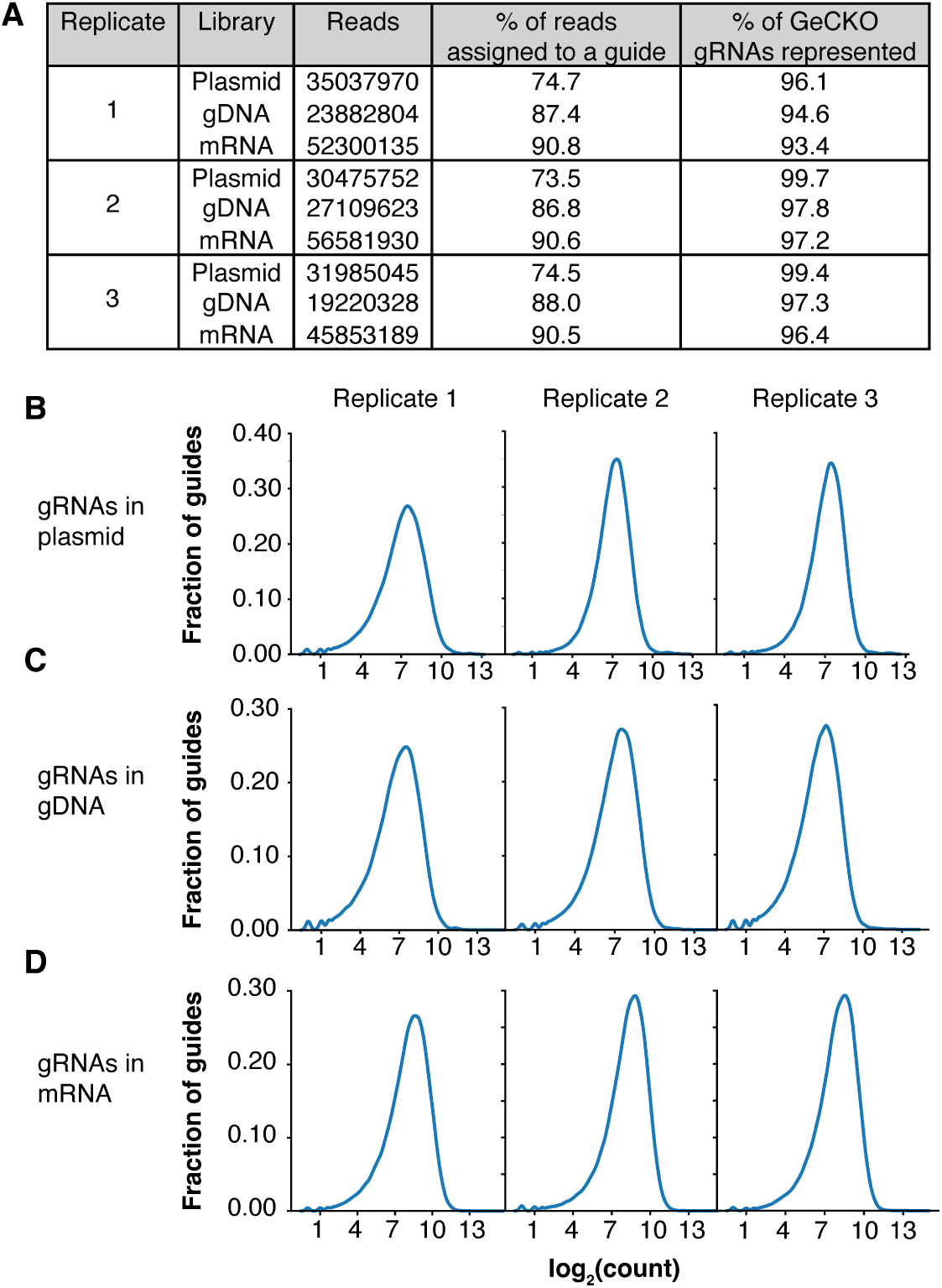
Representation of GeCKO library gRNAs in primary CRITR-seq libraries. (A) Three CRITR-seq libraries were generated by cloning the GeCKO library gRNA sequences into the CRITR-seq vector three independent times. Amplicon sequencing was performed on the region containing the gRNA in the plasmid library, the genomic DNA of the transduced A549 cells, and the mRNA from the transduced cells after 8 hour infection by NS1_mut_. gRNAs were identified for reads matching the CRITR-seq vector for the 10 nucleotides upstream and 13 nucleotides downstream of the gRNA sequence, requiring perfect matching. (B-D) Distribution of gRNAs by count in the plasmid (B), genomic DNA (C), and RNA (D) libraries.

**Supplementary Figure 3.**
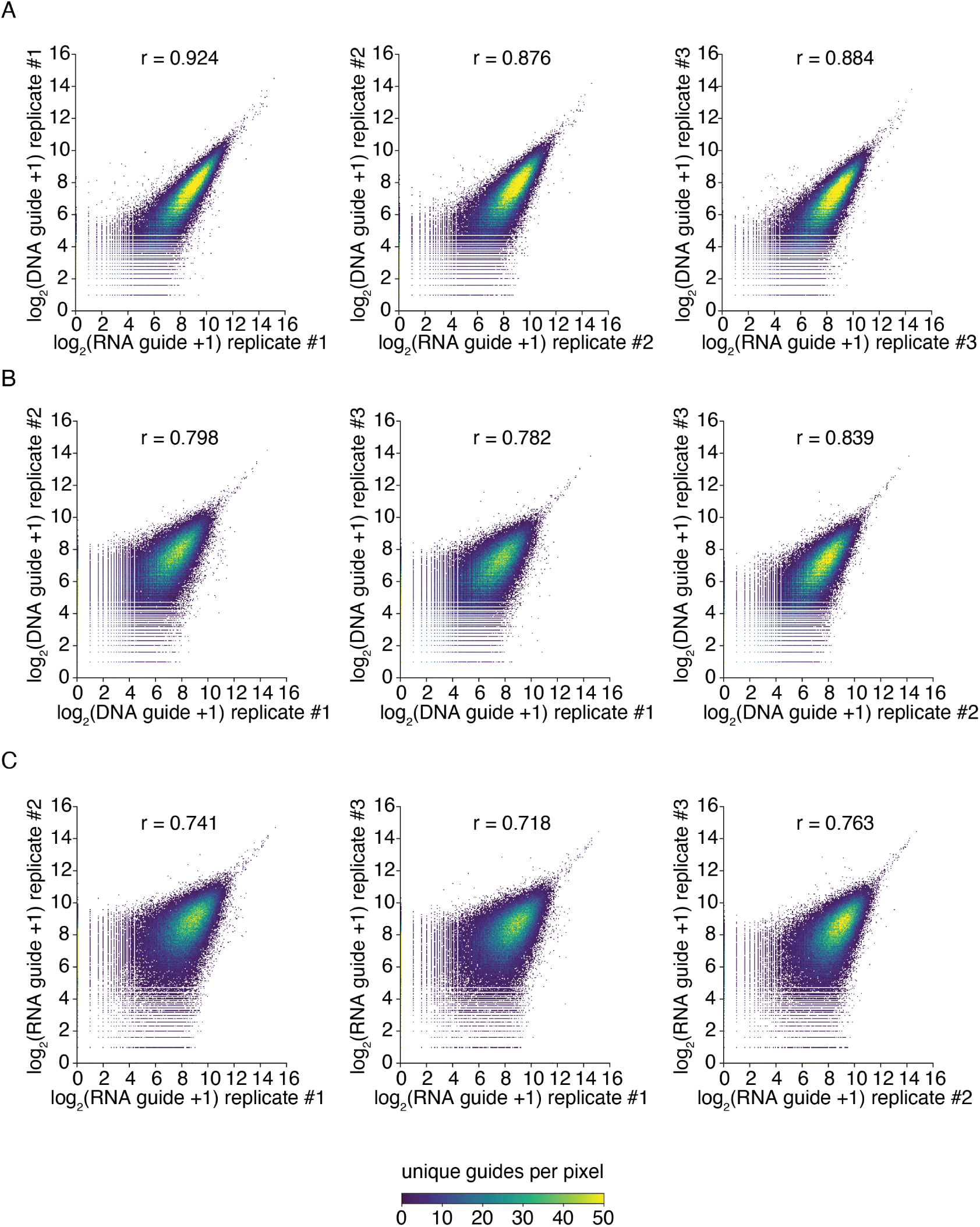
Guide correlation plots between primary CRITR-seq screen replicates. r values are the Pearson correlation coefficient. (A) Guide counts in RNA (x-axis) versus DNA (y-axis). Demonstrates most guides have equivalent effects. (B) Guide count correlations between DNA replicates. Does not correct for differences in initial library, correlations impacted by differences in guide effects on growth and initial distribution as well as other sources of noise. (C) Guide count correlations between RNA replicates. Does not correct for differences in initial library, nor in DNA measurements, correlations impacted by differences in guide effects on growth and initial distribution as well as other sources of noise. Values compare favorably to r values from DNA replicates, indicating little additional noise.

**Supplementary Figure 4.**
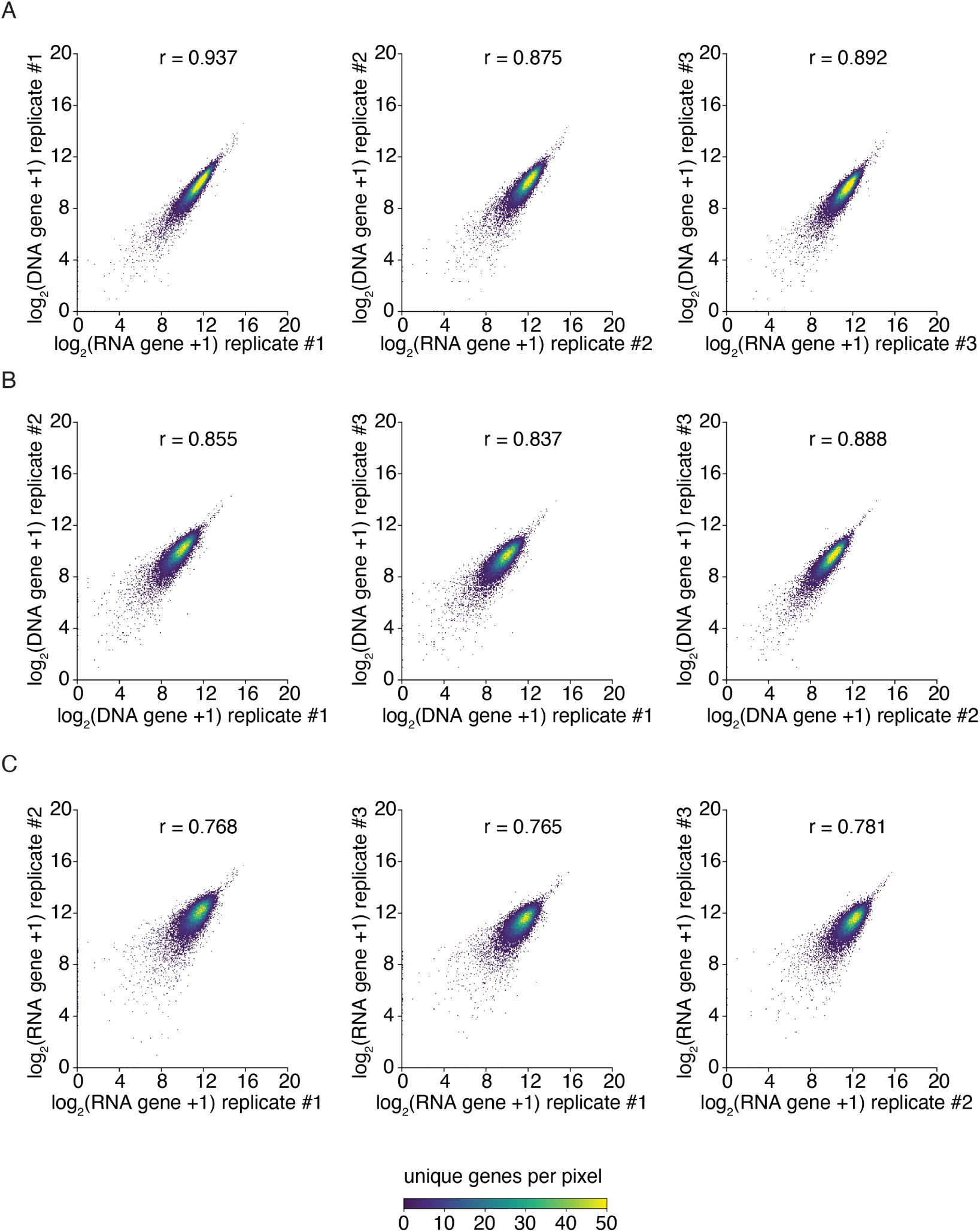
Gene correlation plots between primary CRITR-seq screen replicates. Data plotted as in Fig.S3, but with counts summed for all guides targeting a given gene.

**Supplementary Figure 5.**
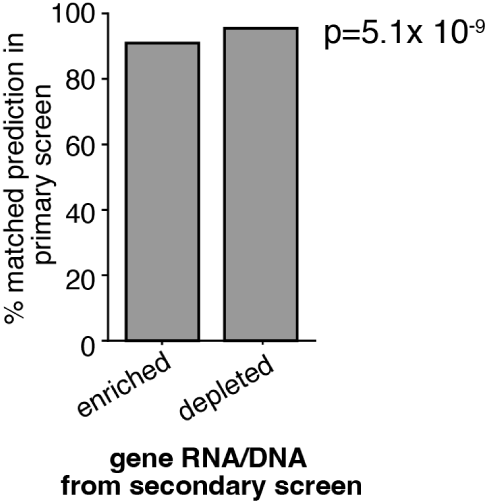
The top 10% of enriched and the top 10% of depleted genes in our smaller screen were compared against their prediction (enriched or depleted) in the larger, primary, screen. p-value Fisher’s exact test. For the smaller screen, inter-replicate correlates at the levels of guides shown in Fig. S6. Full data in Table S2.

**Supplementary Figure 6.**
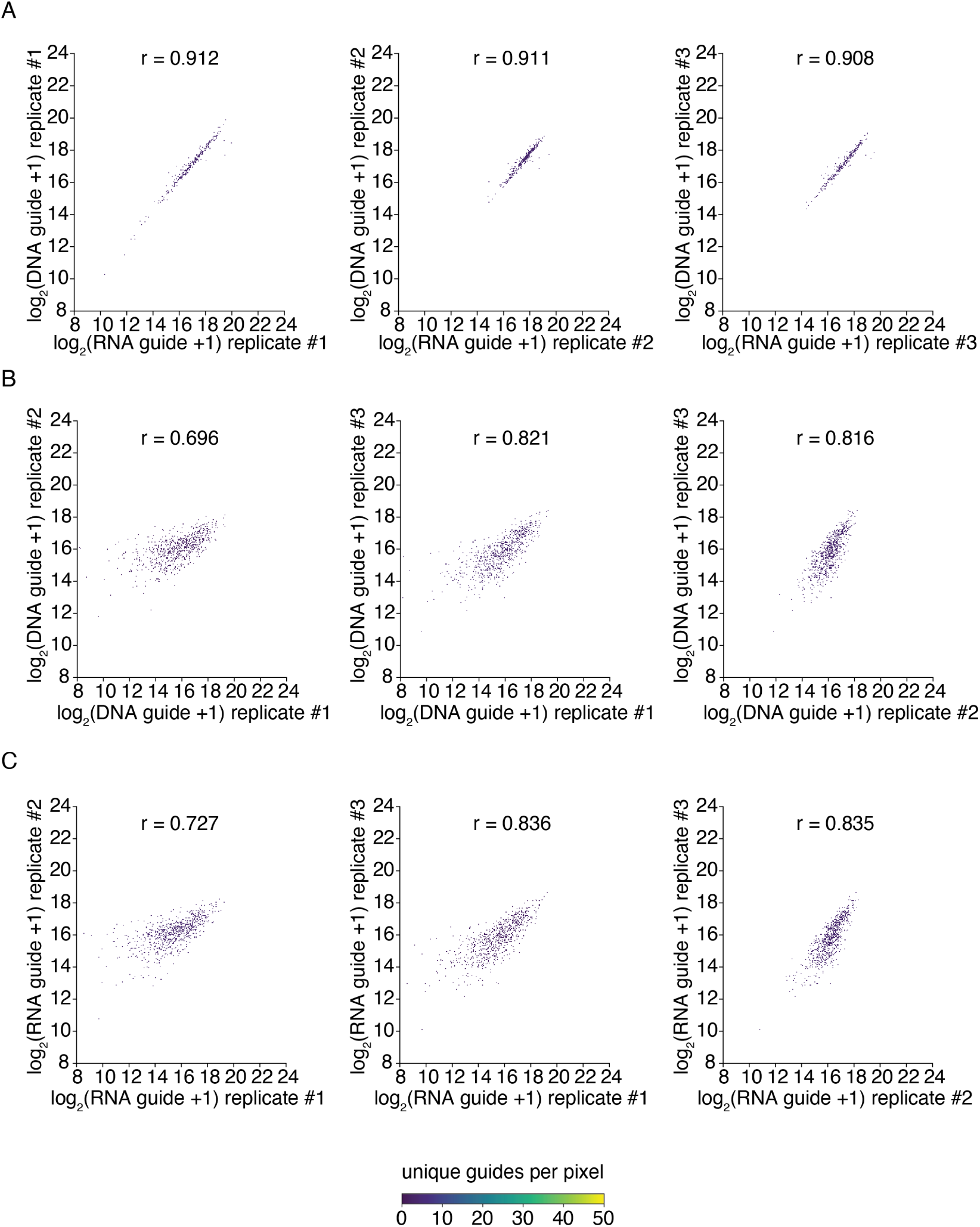
Guide correlation plots between CRITR-seq screen replicates, related to Fig. S5. r values are the Pearson correlation coefficient. (A) Guide counts in RNA (x-axis) versus DNA (y-axis). Demonstrates most guides have equivalent effects. (B) Guide count correlations between DNA replicates. Does not correct for differences in initial library, correlations impacted by differences in guide effects on growth and initial distribution as well as other sources of noise. (C) Guide count correlations between RNA replicates. Does not correct for differences in initial library, nor in DNA measurements, correlations impacted by differences in guide effects on growth and initial distribution as well as other sources of noise. Values compare favorably to r values from DNA replicates, indicating little additional noise.

**Supplementary Figure 7.**
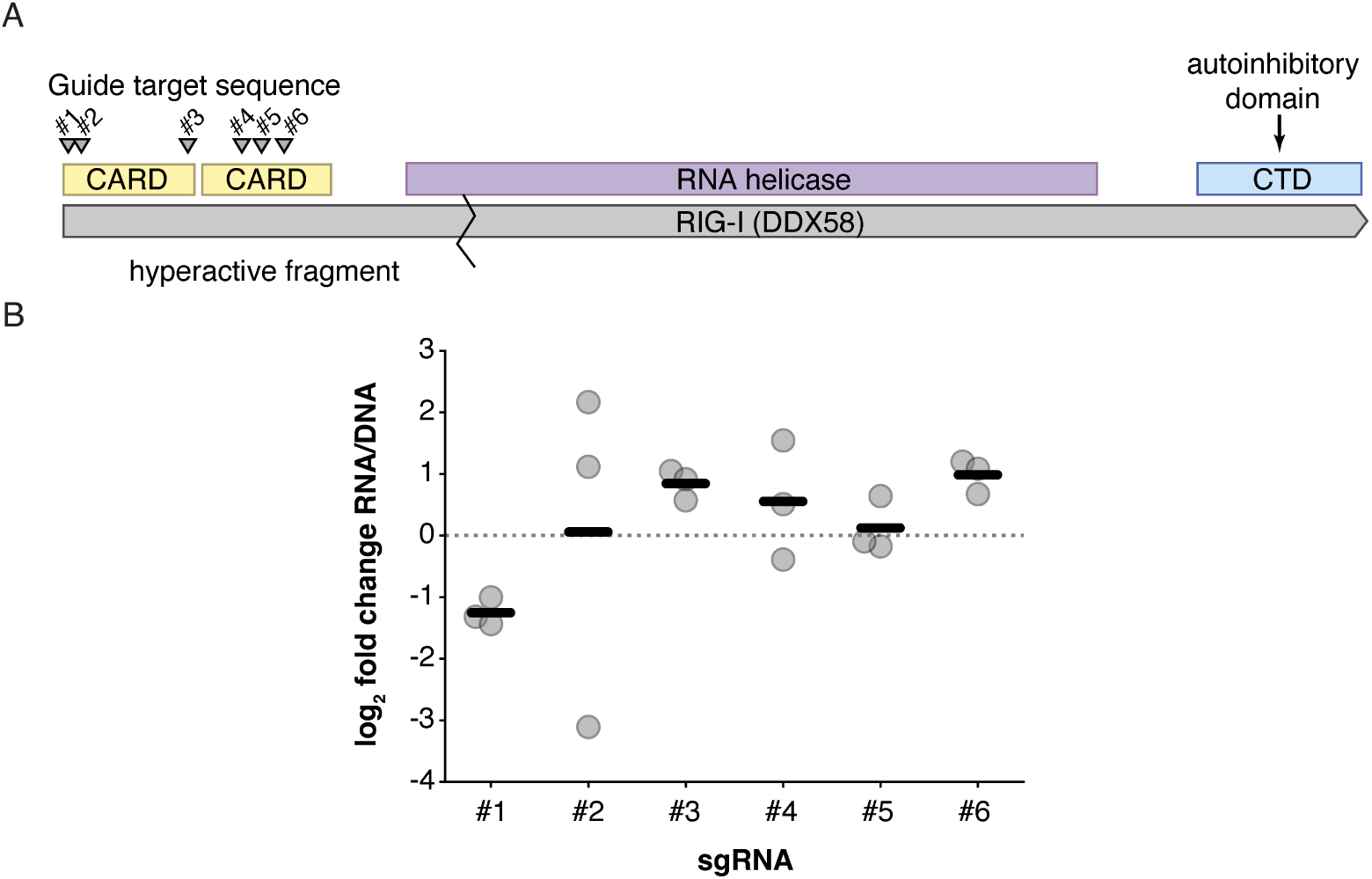
Failure to recover RIG-I as a strong depleted hit may be due to guide distribution. (A) Schematic of RIG-I, consisting of tandem CARD domains, a helicase, and an autoinhibitory C-terminal domain (CTD). A fragment containing just the tandem CARD domains is sufficient to hyperactivate interferon signaling.^90^ While there are no published reports testing a single CARD signaling domain, and as such we do not know whether a single CARD from RIG-I would be weakly positive, positive, or negative, for interferon expression, we note proteins such as MAVS use such an architecture suggesting there may be residual, ligand-independent signaling with a single CARD domain. (B) Measurements across all three replicates for each of the guides indicated in A. Guide #2, with a large spread, is poorly represented across all three libraries, leaving guide #1 as the only guide that would confidently remove both CARD domains.

**Supplementary Figure 8.**
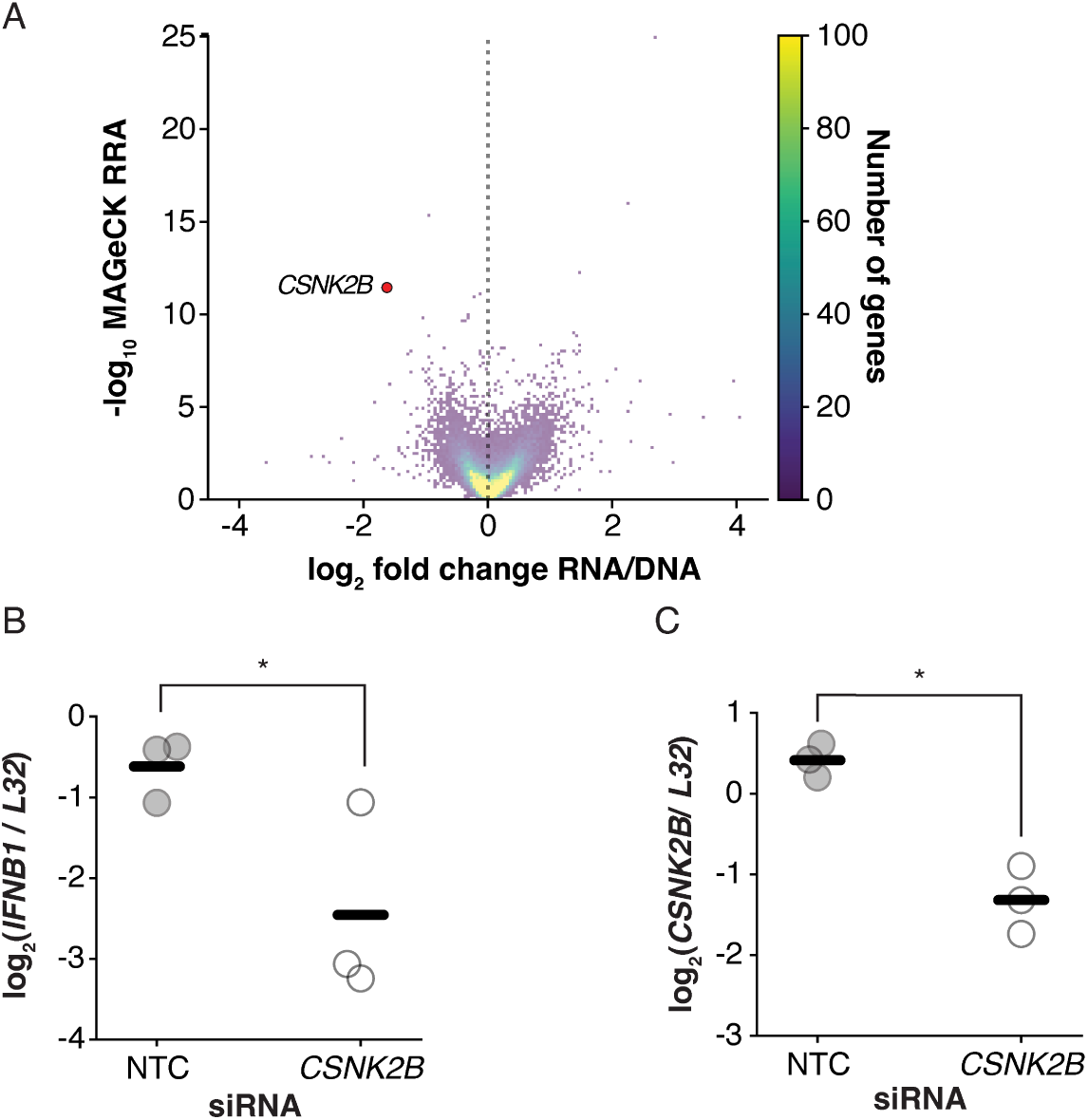
Validation of *CSNK2B* as required for interferon induction during NS1_mut_. Related to Fig. 2 (A) Position of *CSNK2B* in our CRITR-seq dataset as presented in Fig. 2A. (B,C) A549 cells treated with siRNA targeting *CSNK2B* or a non-targeting control for 24h were infected with NS1_mut_ at an MOI of 1 and RNA harvested and analyzed by qPCR at 8h post infection for levels of *IFNB1* and *CSNK2B* transcripts normalized to the housekeeping ribosomal gene *L32*. One-tailed Student’s t-test for a significant drop in either transcript, p<0.05.

**Supplementary Figure 9.**
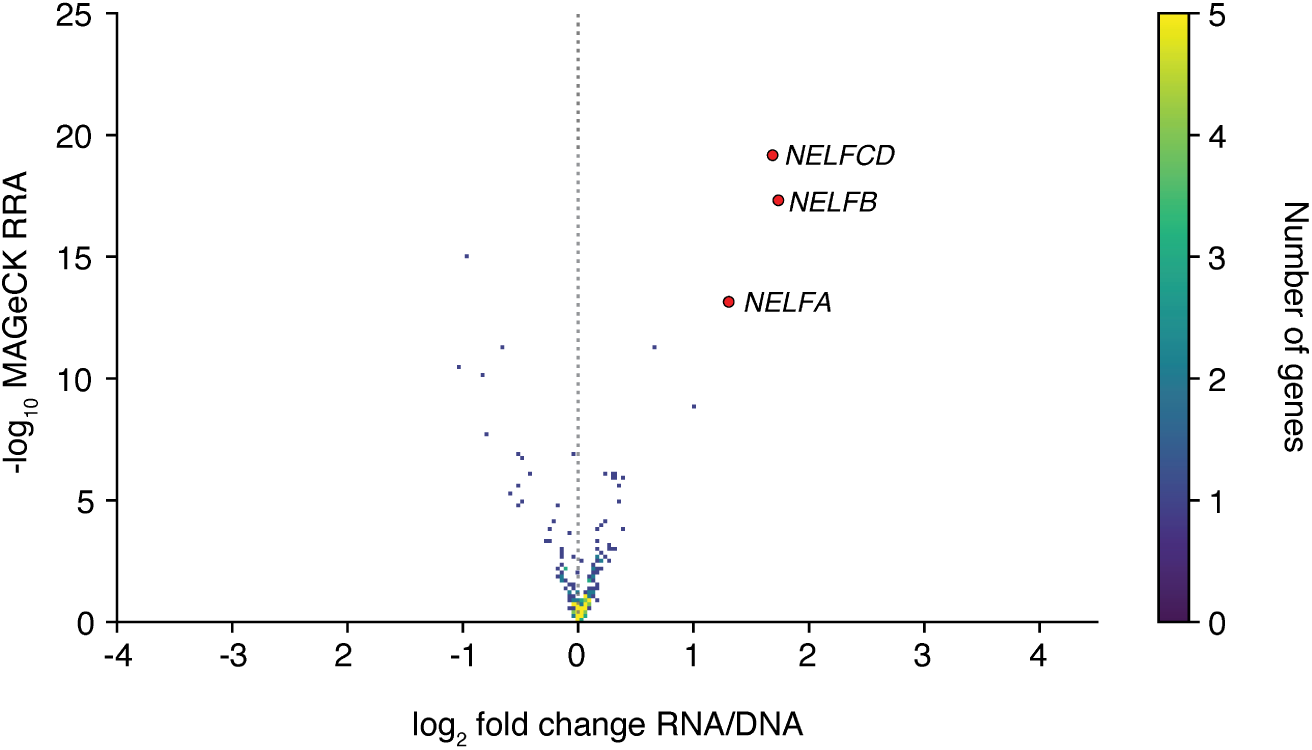
Results from secondary screen as described in Fig S.1, highlighting the top 3 enriched genes by MAGeCK analysis. Full data presented in Table S2.

**Supplementary Figure 10.**
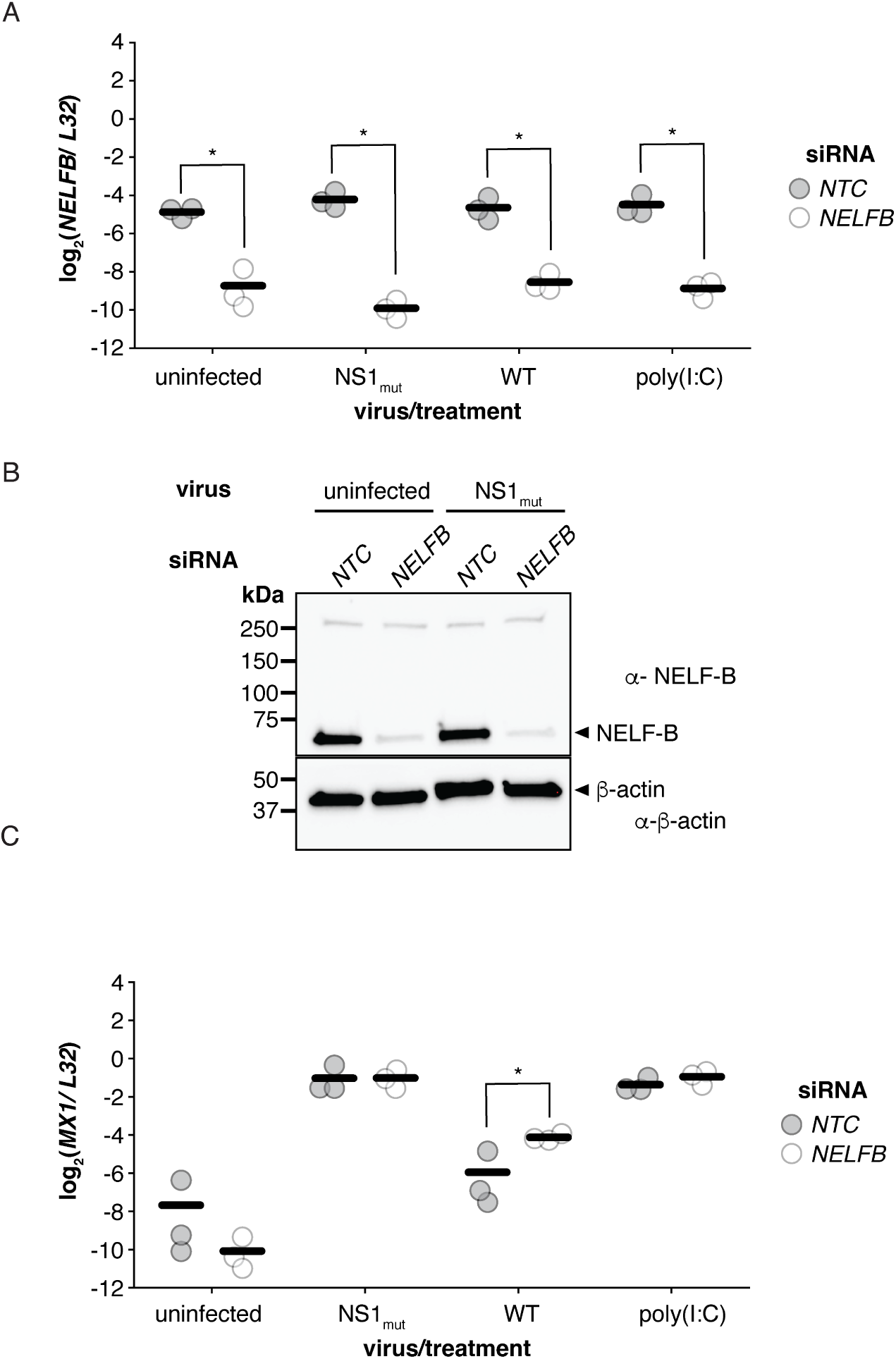
Validation experiments for Fig. 3C. (A) qPCR measurements of *NELFB* normalized to *L32* from samples in Fig.3C. Asterisks indicate significant depletion, one-tailed Student’s T-test with Benjamini-Hochberg multiple hypothesis, p<0.05, n=3. (B)Western blot analysis of lysate from indicated samples contemporaneously generated as those in Fig. 3C. Lysates were probed for NELFB protein and the loading control β-actin. Bands matching the predicted size of each protein highlighted. (C) Analysis as in (A) measuring the interferon-stimualted gene *MX1*. Asterisks indicate significant difference, one-tailed Student’s t-test for an increase upon *NELFB* silencing with Benjamini-Hochberg multiple hypothesis correction, p<0.05, n=3. No significant differences in ISG expression oberved upon *NELFB* silencing without flu infection. Lack of differences in NS1_mut_ experiments may reflect a saturated interferon signaling pathway, as we still observe a difference during a wild-type infection where levels of interferon expression are lower.

**Supplementary Figure 11.**
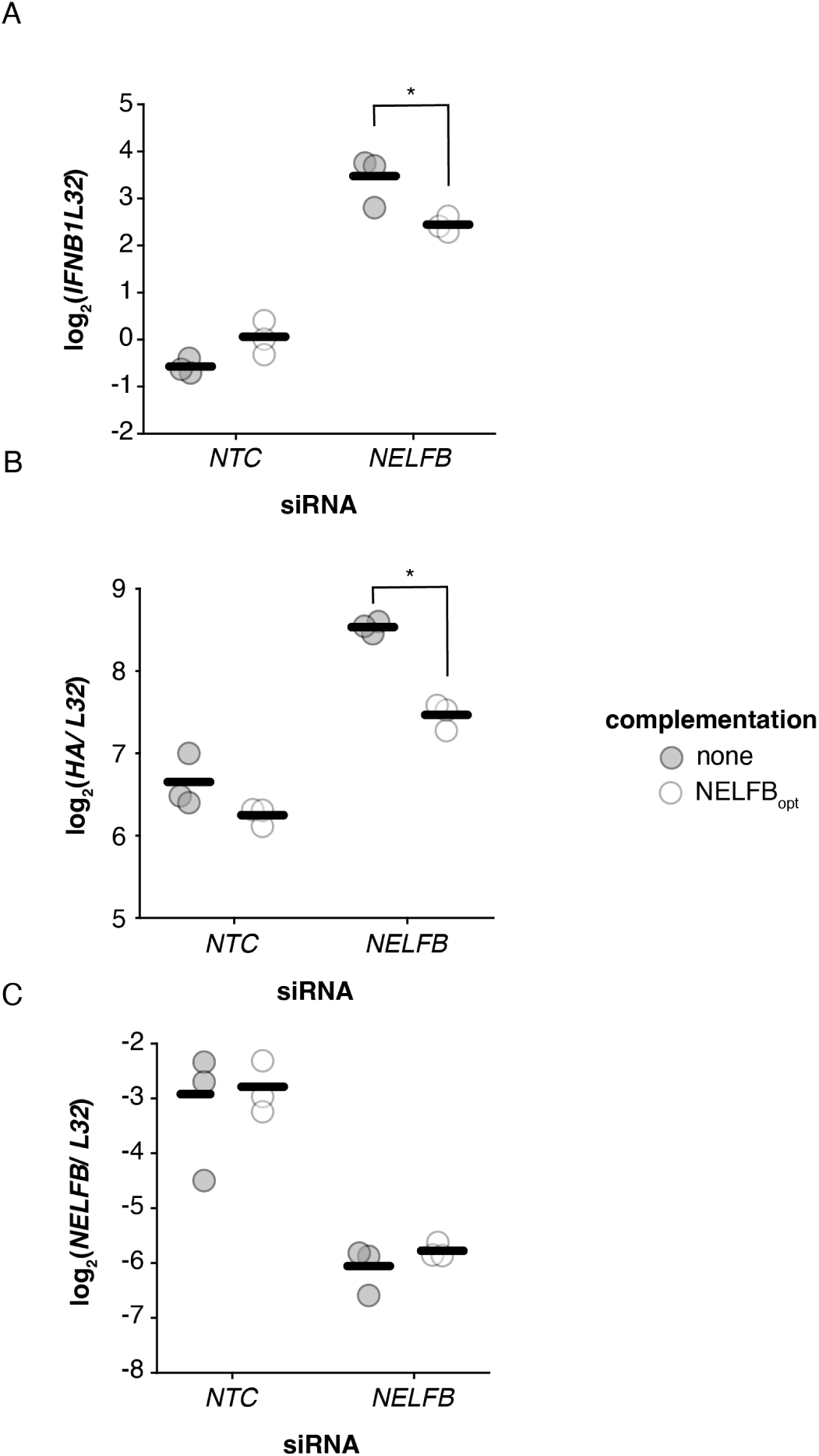
Overexpression of silencing-resistant *NELFB* rescues phenotypes observed in this study, related to Figs. 3, 4 (A-C) Cells were silenced as in Fig. 3C. 19.5h prior to infection, cells were either transduced with a lentivirus expressing a codon-optimized silencing-resistant *NELFB* or treated identically with no transduction. Cells were then infected as in Fig. 3C, and RNA harvested 8h post-infection and analyzed by qPCR for the indicated transcripts. Asterisks represent significant decreases from the no-complementation control, one-tailed Student’s t-test, with Benjamini-Hochberg multiple hypothesis correction, n=3, p<0.05. Differences were only observed in *NELFB* –silenced cells, consistent with the proposed role of NELFB acting in concert as part of the NELF complex such that an excess of NELFB would not have an effect. Silencing of endogenous *NELFB* was unimpacted by our transduction. The expression construct we used for silencing-resistant NELF is not captured by this qPCR, allowing us to measure endogenous levels alone and confirm our manipulations did not interfere with silencing.

**Supplementary Figure 12.**
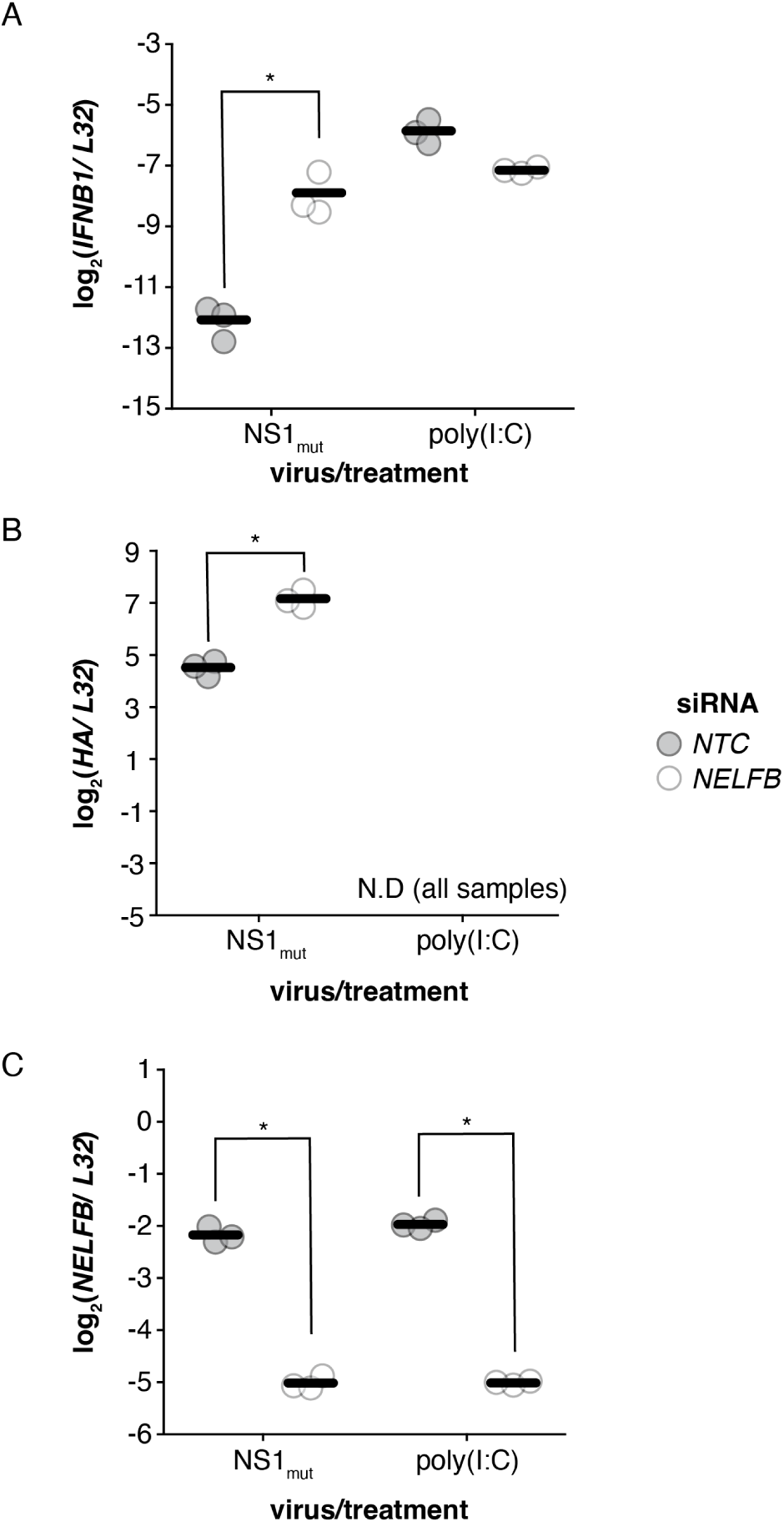
Silencing of *NELFB* in 293T cells increases interferon transcription in response to flu and not poly(I:C) and expression of flu as seen in A549 cells. Related to Figs. 3,4. (A-C) 293T cells were treated with siRNA for 9 days and then infected with NS1_mut_, left untreated, or transfected with 50ng poly(I:C). RNA was harvested 8h post treatment and analyzed for the indicated transcripts by qPCR. Asterisks indicate significant increases (A,B) or decreases (C) between control and *NELFB* silencing, one-tailed Student’s t-test with Benjamini-Hochberg multiple hypothesis correction, n=3, p<0.05.

**Supplementary Figure 13.**
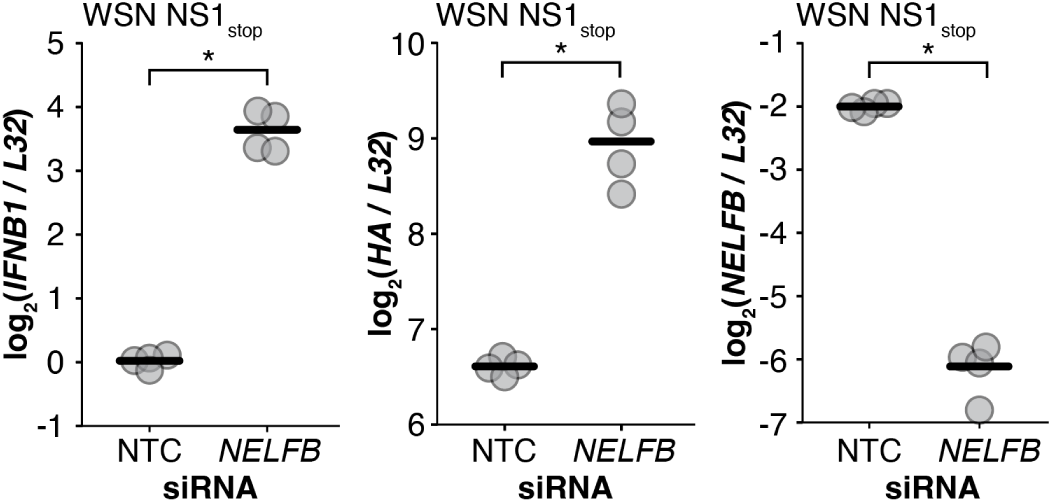
Effects of *NELFB* knockdown on WSN NS1_stop_ IAV, related to Fig. 3, 4. The NS1_mut_ virus used elsewhere in this study has the WSN genome segments 1-7 and PR8 segment 8 with early stop mutations in the NS1 open reading frame that do not affect the coding sequence of NS2. This virus was used for the CRITR-seq screen and most follow-up experiments because it grows to higher titers than the NS1_stop_ virus in the fully WSN background. To validate the effects of *NELFB* are not particular to the NS1_mut_ virus with mixed genetic background, we treated cells with either non-targeting control or *NELFB* siRNA for 9 days, followed by WSN NS1_stop_ infection at an MOI of 1. RNA was harvested 8 hours post infection for qPCR analysis. Biological replicates are shown as individual data points, with lines representing the means. Two-tailed t-tests were performed to compare *NELFB* knockdown with the non-targeting control in each condition. n=4, with two replicates each of two biological replicate viral populations. * indicates p<0.05.

**Supplementary Figure 14.**
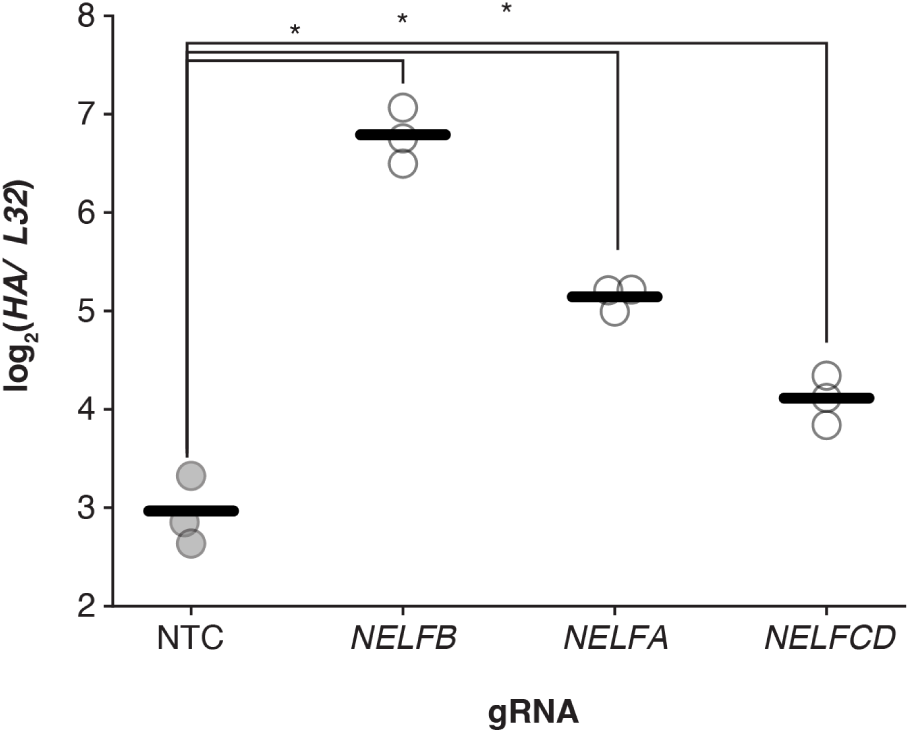
Cas9 knockouts of *NELF* genes increase influenza RNA. RNA from Fig. 3B was analyzed for flu HA levels. Asterisks indicate significance, two-tailed Student’s t-test with Benjamini-Hochberg multiple hypothesis correction, n=3, p<0.05.

**Supplementary Figure 15.**
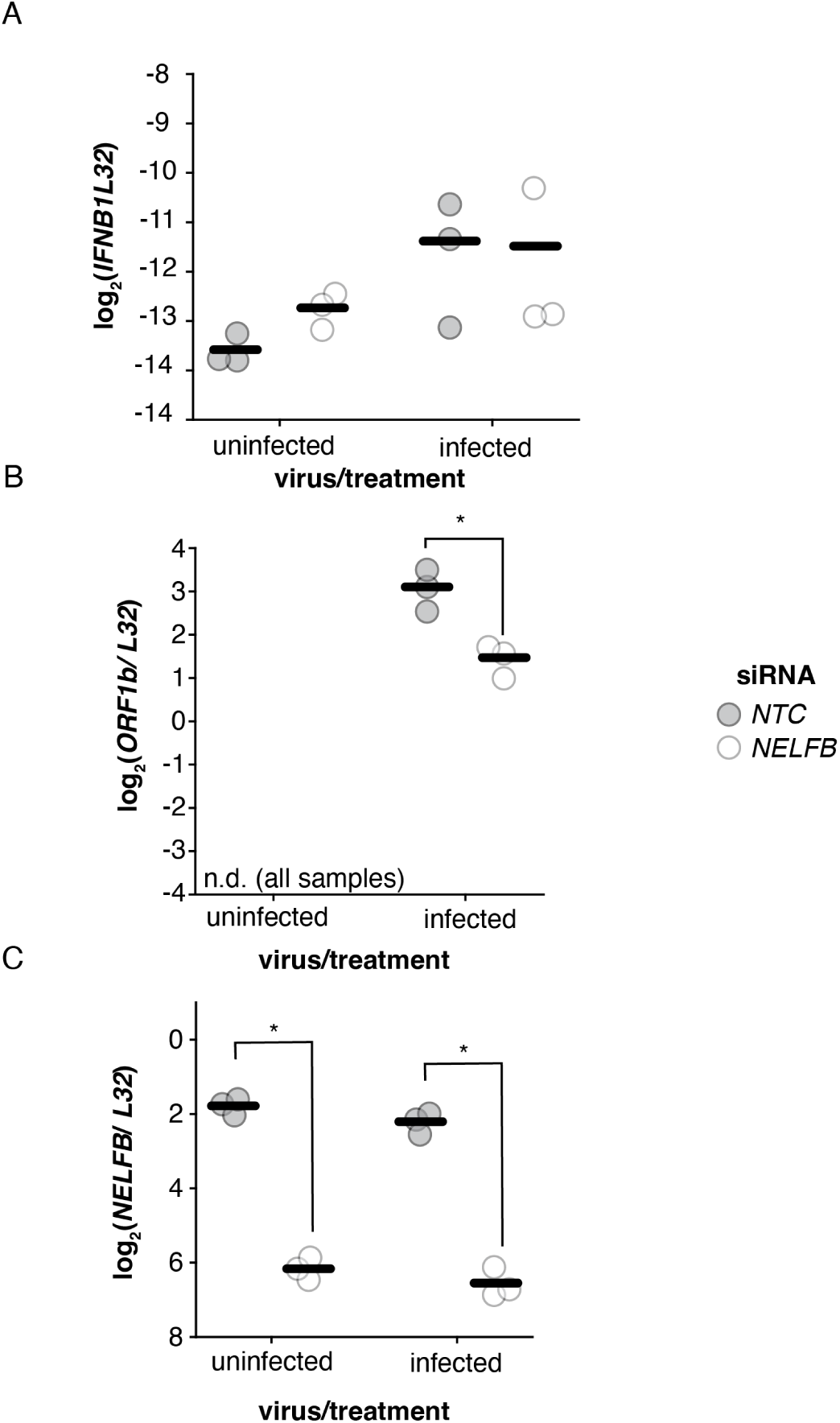
Silencing *NELF* does not increase coronavirus interferon production or replication. (A) A549 cells were treated with siRNA for 9 days. Cells were infected, or not, with coronavirus OC43 at an MOI of 1 for 21h and RNA was harvested and analyzed by qPCR. Each treatment analyzed by two-tailed Student’s t-test, no significance detected between non-treatment control and *NELFB* silencing, p<0.05, n=3 with Benjamini-Hochberg multiple hypothesis correction. (B) RNA analyzed from A with primers against the coronavirus *ORF1b*. Significance analyzed as in A. (C) Confirmation of silencing from A and B.

**Supplementary Figure 16.**
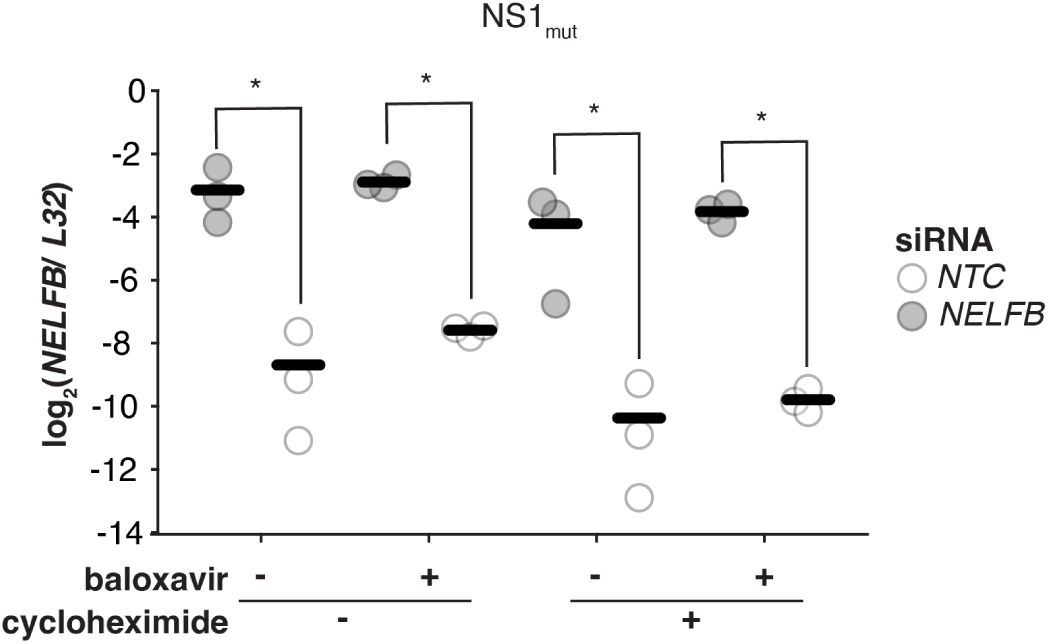
qPCR measuring *NELFB* silencing in Fig. 4B. Asterisks indicate significant decreases in *NELFB* transcripts when corrected for the housekeeping control *L32*, one-tailed Student’s t-test with Benjamini-Hochberg multiple hypothesis correction, p<0.05, n=3.

**Supplementary Figure 17.**
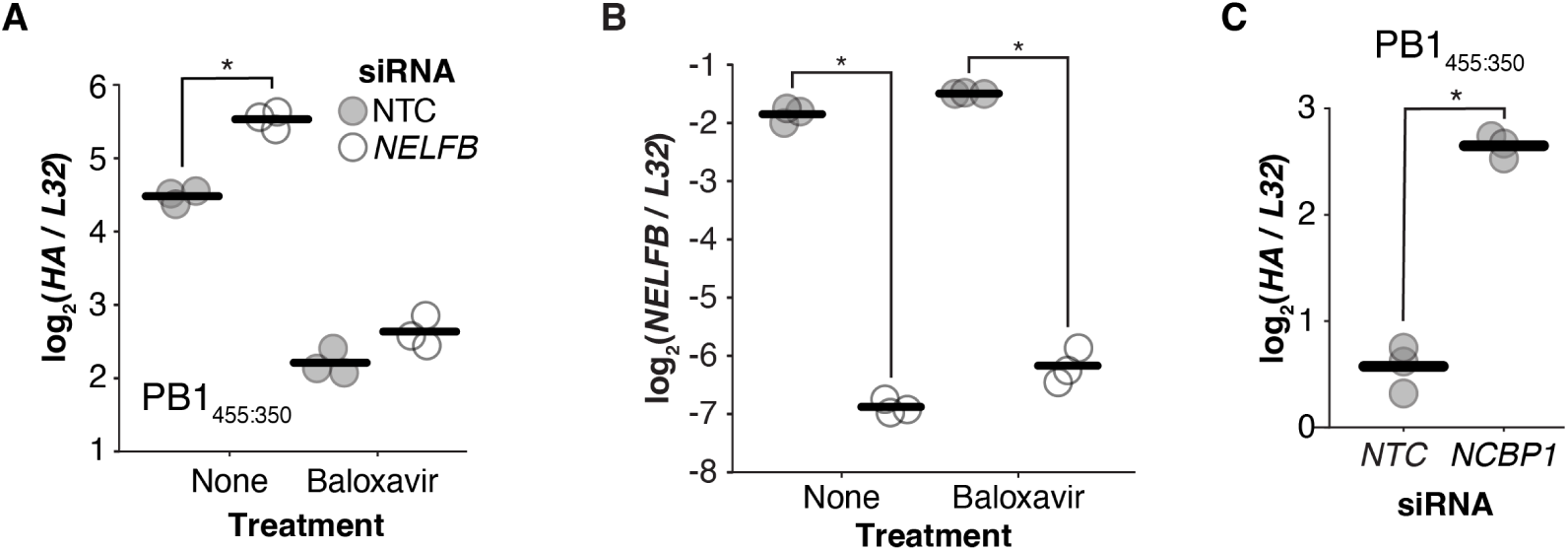
Influence of *NELFB* and *NCBP1* silencing on primary transcription. (A)A549 cells were treated with siRNA for 9 days. Cells were infected with PB1_455:350_ at an infectious MOI of 1, with or without 100 nM baloxaviric acid added at time of infection. RNA was harvested 8 hours post infection for analysis by qPCR. (B) Validation of *NELFB* silencing from A. (C) A549 cells were treated with siRNA for 4 days and infected with PB1_455:350_ at an infectious MOI of 1. RNA was harvested 8 hours post infection for analysis by qPCR.

**Supplementary Figure 18.**
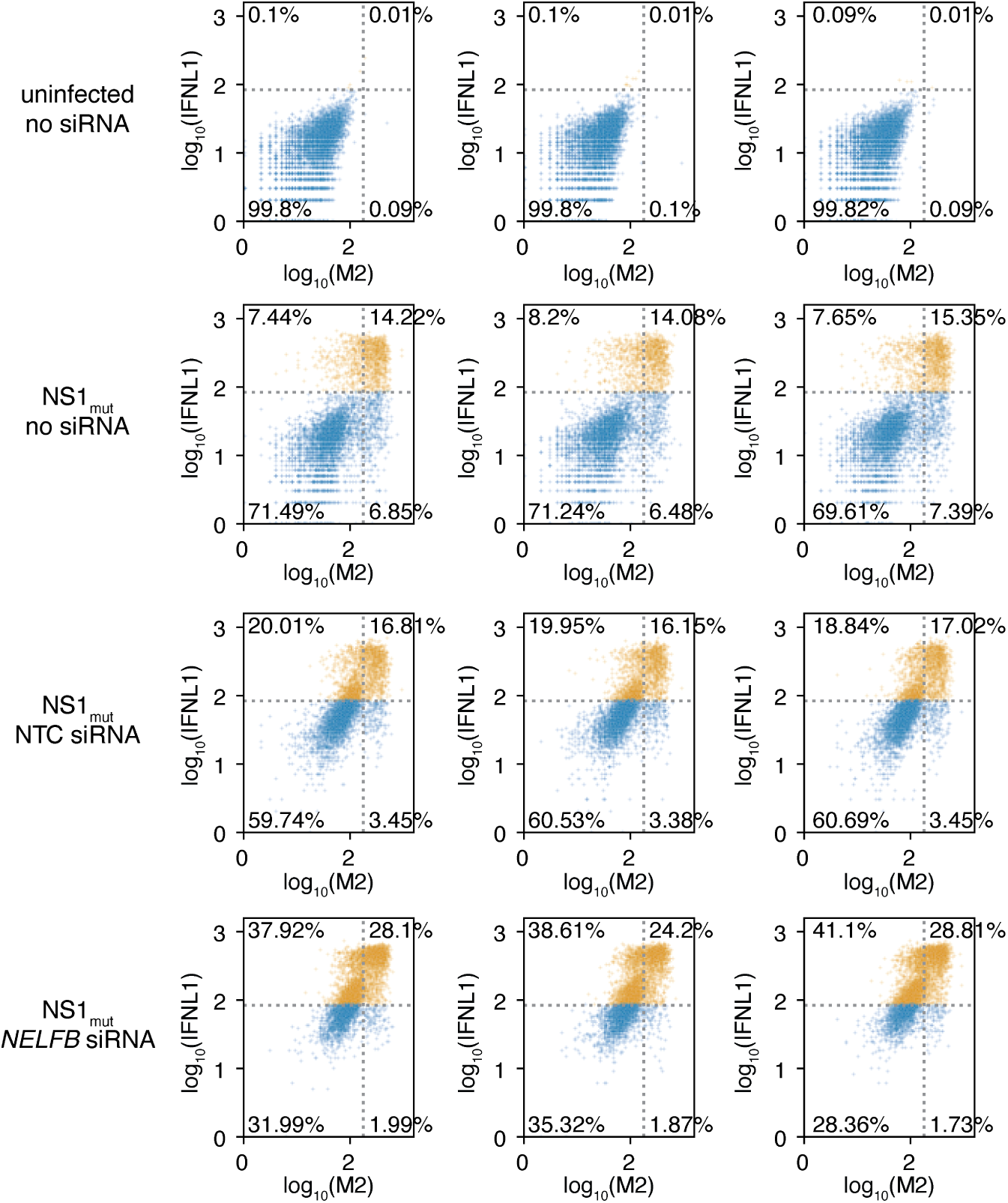
Full flow cytometry data, related to Figs. 4, 5. A549 *IFNL1* reporter cells were treated for 9 days with siRNA and infected, or not, with NS1_mut_ at a genome-corrected MOI of 1. 13 hours post infection, cells were stained for the viral protein M2 and fixed for flow cytometry. Individual replicates shown. The threshold for IFNL1^+^ or M2^+^ cells was set at the fluorescence level for which an average of 0.1% of uninfected cells would be called as positive events. Interferon-positive events colored in orange. For visualization, data was subsetted to 5000 events to show equivalent numbers between conditions.

**Supplementary Figure 19.**
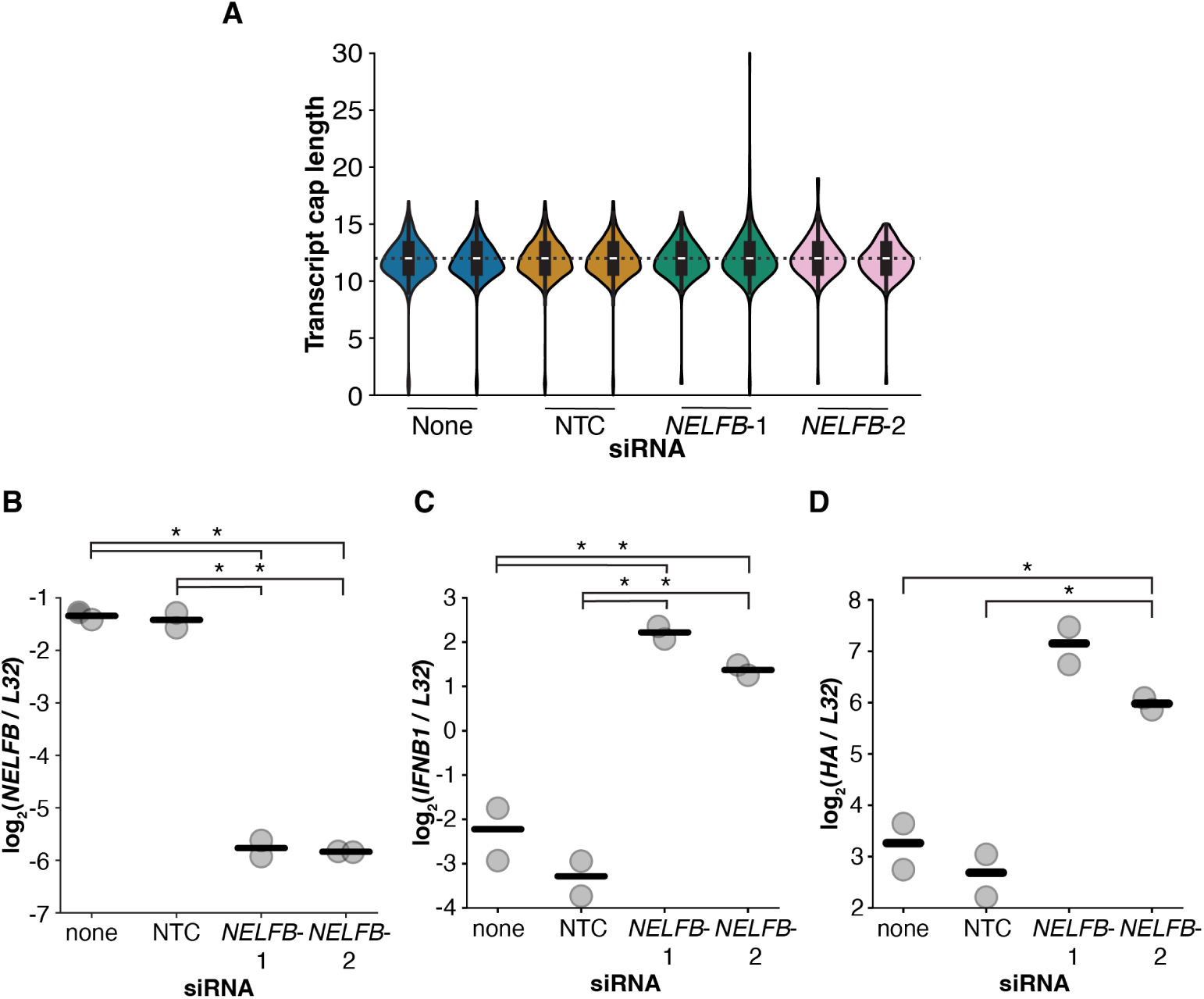
*NELFB* silencing does not appear to influence length of host-derived influenza cap sequences. (A)A549 cells were treated with either of 2 different *NELFB* –targeting siRNAs, a non-targeting control, or no siRNA, with 2 biological replicates per treatment. After 9 days of siRNA treatment, cells were infected with NS1_mut_ at an MOI of 2. RNA was harvested 8 hours post infection, and 5’ RACE was performed on polyadenylated mRNAs, followed by sequencing. Cap-snatched sequence length refers to the number of nucleotides between the template switch oligo sequence and the +1 position of the flu mRNA sequence. Violin plots contain box plots for each sample, and the median of all samples is represented by the gray dotted line. Violin plot whiskers extend to the most extreme points in the dataset, excluding the top 2% of lengths. (B-D)qPCR of RNA used in A demonstrating decreases in *NELFB*, increases in interferon, and increases in flu transcription, consistent with all other experiments. Biological replicates are shown as individual data points, with lines representing the means. Two-tailed t-tests were performed to compare each sample with the untreated condition and with the non-targeting control, n=2. * indicates p<0.05 after Benjamini-Hochberg multiple hypothesis correction.

**Supplementary Figure 20.**
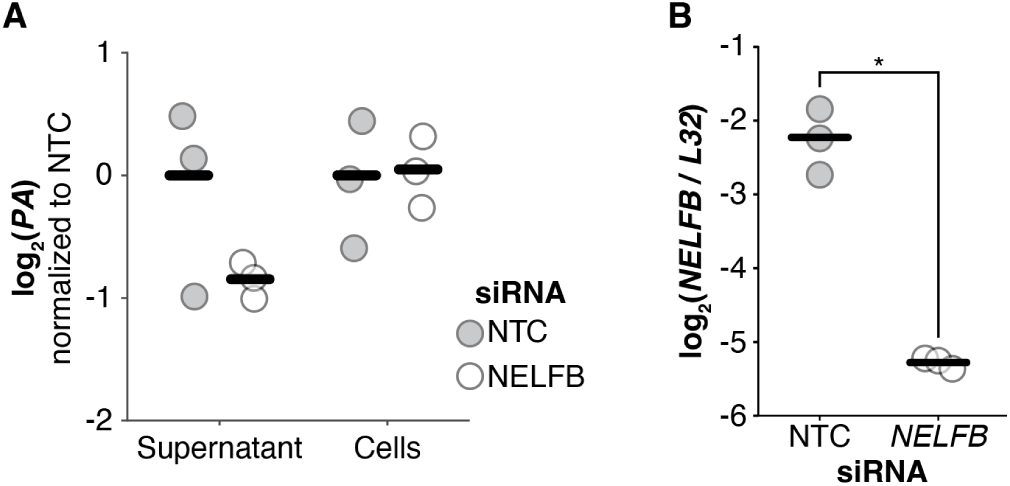
*NELFB* knockdown does not increase viral titer. (A-B) Same experiment as in Fig. 4E, with different RNAs measured by qPCR. A549 cells were treated with siRNA for 9 days before infection with WT WSN at an infectious MOI of 5. Media was replaced with fresh IGM 2 hours post infection. 14 hours post infection, viral supernatant was collected and cells were lysed for RNA extraction of both. Reverse transcription was performed using universal influenza primers for the RNA from the supernatant, and random hexamer primers for the RNA from the cell lysate. Biological replicates are shown as individual data points, with lines representing the means. Two-tailed t-tests were performed to compare targetign siRNA with the non-targeting control, n=3. Benjamini-Hochberg multiple hypothesis correction was performed for panel A. * indicates p<0.05.

**Supplementary Figure 21.**
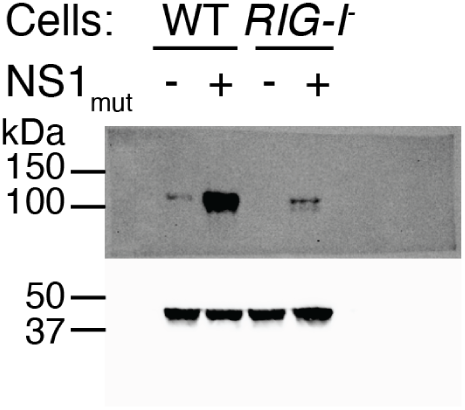
RIG-I protein expression in clonal *RIG-I* knockout A549 cell line. We transfected A549 cells with Cas9 ribonucleoprotein using a gRNA targeting *RIG-I*. After performing dilution cloning to isolate single cells, the clonal-derived line was infected with NS1_mut_ at an MOI of 5 for 24 hours. Infected and uninfected A549 or A549 RIG-I knockout cells were lysed for SDS-PAGE and Western blot.

**Supplementary Figure 22.**
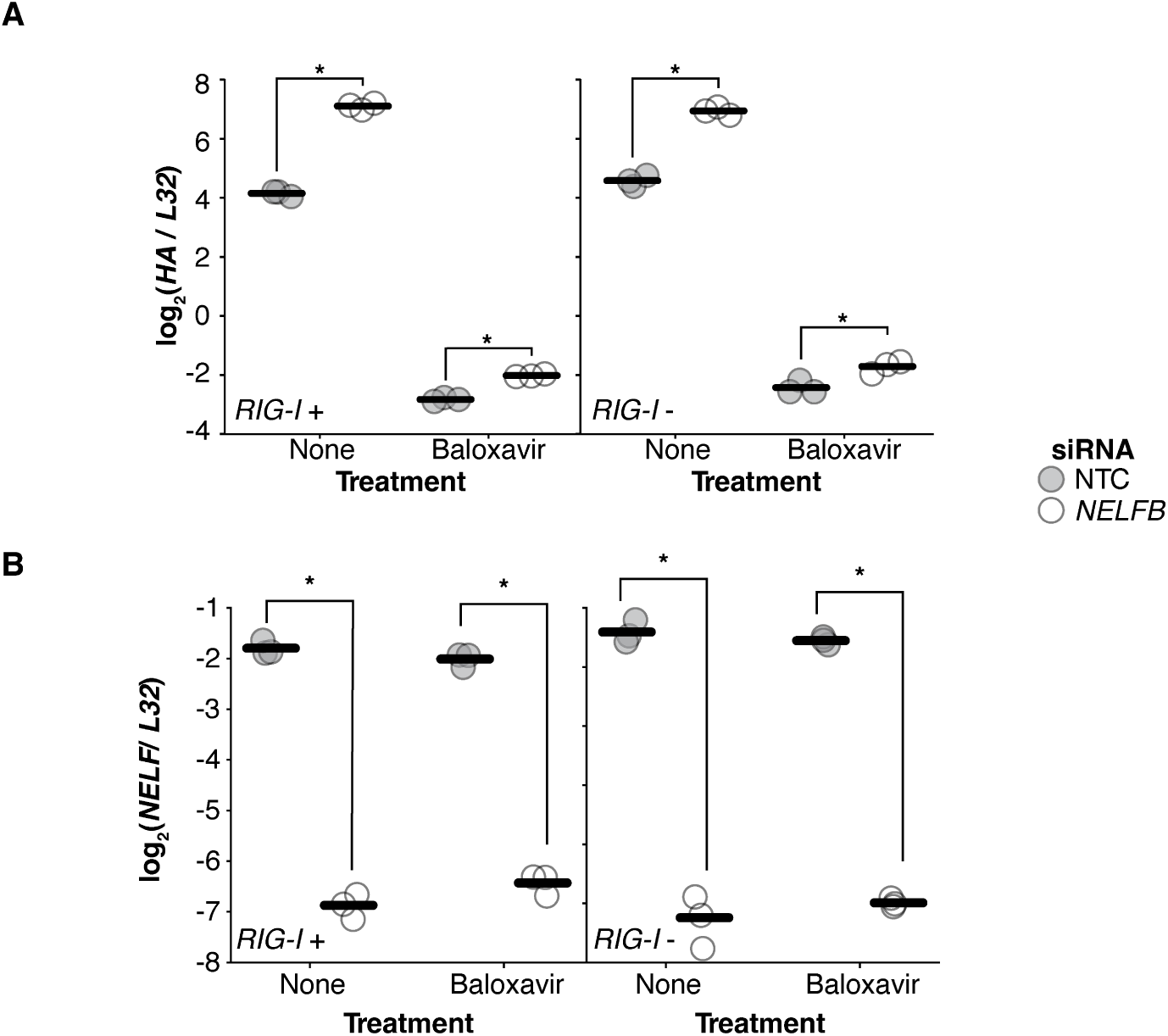
*RIG-I* knockout does not affect *HA* RNA levels. (A)Same experiment as in Figs. 5A and B, but measuring *HA* levels by qPCR. A *RIG-I* knockout cell line derived from a single cell clone (Fig. S21) was treated with siRNA for 9 days. Cells were infected with NS1_mut_ at a genome-corrected MOI of 1, with or without 100 nM baloxaviric acid added at time of infection. RNA was harvested 8 hours post infection for qPCR analysis. Points represent biological replicates, with lines indicating the means. Two-tailed t-tests were performed comparing the NELFB siRNA samples with the non-targeting control for each treatment condition, n=3. * indicates p<0.05 after Benjamini-Hochberg multiple hypothesis correction. (B) Confirmation of *NELFB* silencing, analyzed as in (A).

**Supplementary Figure 23.**
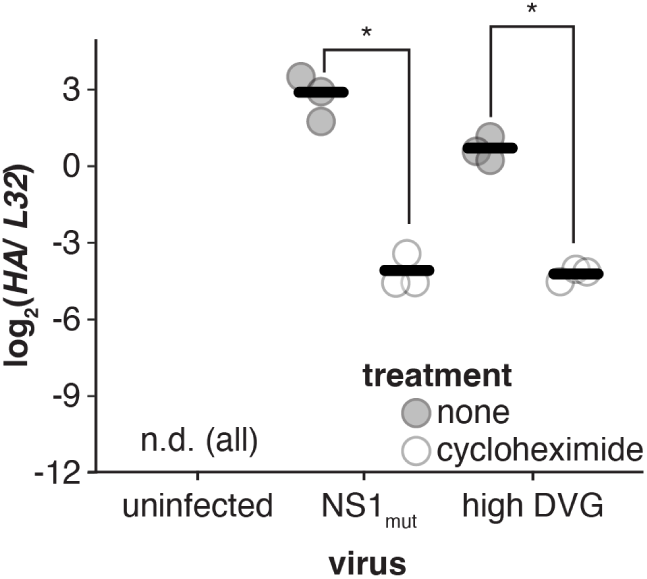
*HA* RNA levels from Fig 6C. Cycloheximide restricts viral replication, and similar levels of HA are observed between flu variants under cycloheximide treatment, indicating similar doses of virus were used. Infectious units cannot be used to appropriately calibrate infections with DelVG populations. Points represent biological replicates, with lines indicating the means. Two-tailed t-tests were performed to identifiy significance upon cycloheximide treatment, n=3. * indicates p<0.05 after Benjamini-Hochberg multiple hypothesis correction.

**Supplementary Figure 24.**
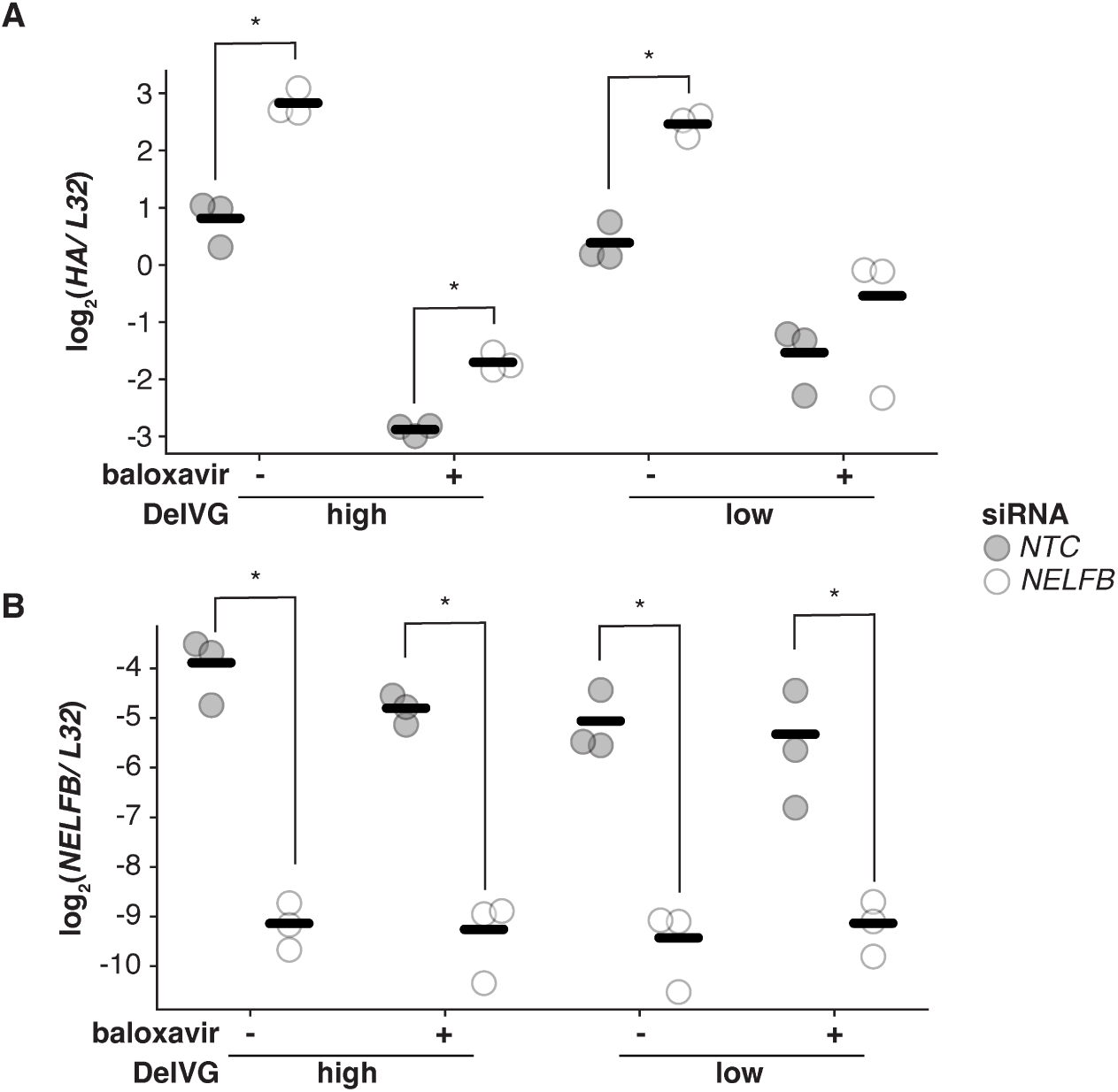
Analysis of RNA from Fig. 6D for *HA* (A) or *NELFB* (B). Points represent biological replicates, with lines indicating the means. Two-tailed t-tests were performed to identifiy significance upon *NELFB* silencing, n=3. * indicates p<0.05 after Benjamini-Hochberg multiple hypothesis correction.

**Supplementary Figure 25.**
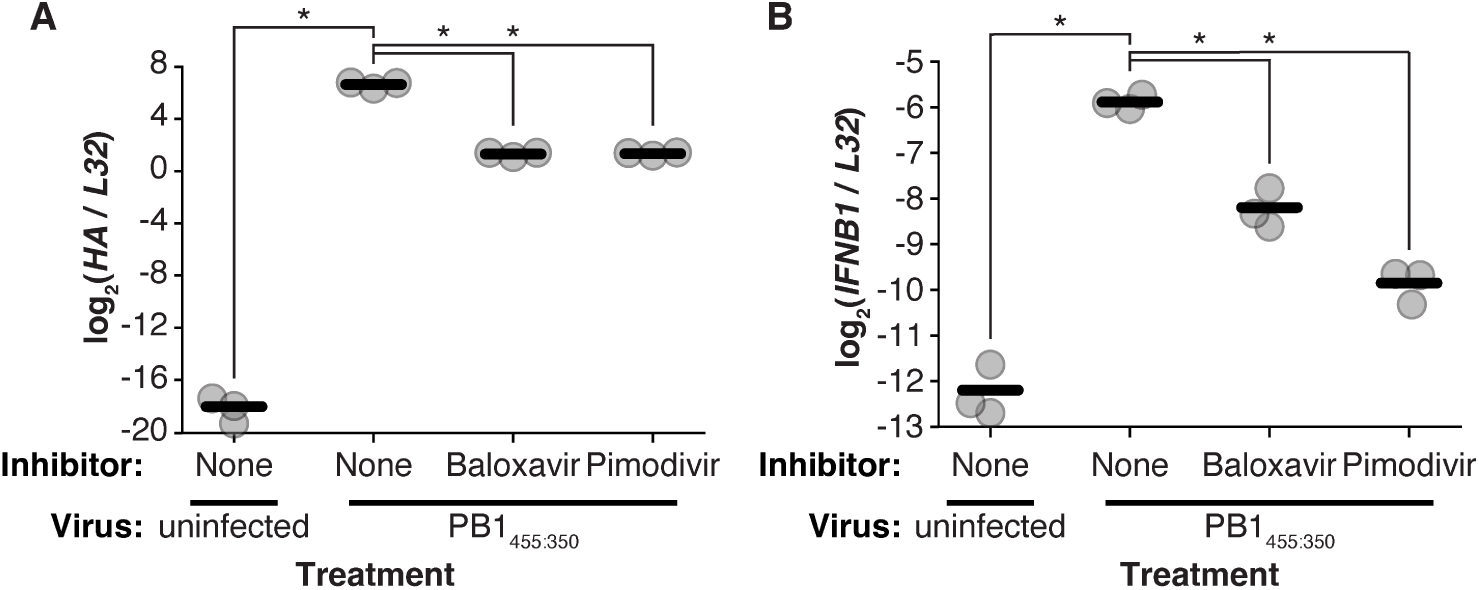
(A-B) A549 cells were infected, or not, with PB1_455:350_, at an infectious MOI of 1, with or without 100 nM baloxaviric acid or 100 nM pimodivir, a compound that inhibits transcription by preventing flu binding to host cap structures, added 2 hours prior to infection. RNA was harvested 14 hours post infection for qPCR analysis. Points represent biological replicates, with lines indicating the means. For A and B, two-tailed t-tests were performed comparing each treatment with the uninfected sample and with the infected sample without inhibitors, n=3.

**Supplementary Table 1. Results from primary screen**

Results from the primary CRITR-seq screen

**Supplementary Table 2. Results from secondary screen**

Results from the secondary CRITR-seq screen

**Supplementary Table 3. Proviral genes**

List of proviral genes as previously identified in Li *et al*. 2020^42^, genes involved in early viral processes indicated, and genes used to assess RIG-I signaling, core genes indicated.

**Supplementary Table 4. crRNA sequences**

crRNA sequences used for editing in this paper.

**Supplementary Table 5. Primers**

Primer sequences used in this paper.

**Supplementary Table 6. siRNAs**

siRNAs used in this paper.

**Supplementary Table 7. sgRNA secondary screen**

sgRNAs cloned and analyzed in our secondary screen.

**Supplementary File 1. Cas9 lentiviral vector**

Sequence of lentiviral vector used to introduce Cas9 into A549 cells.

**Supplementary File 2. Flu genome sequence**

Sequence of flu strains used in this study

**Supplementary File 3. CRITR-seq vector**

Type III interferon CRITR-seq vector generated and used in this study.

**Supplementary File 4. NELF-B complementation vector**

*NELFB* complementation construct. Codon optimization leads to mismatches in region targeted for silencing.

